# The Swiss Army Knife of Alginate Manipulation – A Gut Bacterium Alginate Lyase with Diverse Catalytic Activities

**DOI:** 10.1101/2024.12.11.627697

**Authors:** Tobias Tandrup, José Pablo Rivas-Fernández, Mikkel Madsen, Mette E. Rønne, Agnes B. Petersen, Leesa J. Klau, Anne Tøndervik, Casper Wilkens, Finn L. Aachmann, Carme Rovira, Birte Svensson

**Affiliations:** Enzyme and Protein Chemistry, Department of Biotechnology and Biomedicine, Technical University of Denmark, DK-2800 Kongens Lyngby, Denmark; Departament de Química Inorgánica i Orgánica (Secció de Química Orgánica) and Institut de Química Teorica i Computacional (IQTCUB), Universitat de Barcelona, 08028 Barcelona, Spain; Norwegian Biopolymer Laboratory (NOBIPOL), Department of Biotechnology and Food Science, NTNU Norwegian University of Science and Technology, NO-7491 Trondheim, Norway; Department of Biotechnology and Nanomedicine, SINTEF AS, NO-7491 Trondheim, Norway; Structural Enzymology, Department of Biotechnology and Biomedicine, Technical University of Denmark, DK-2800 Kongens Lyngby, Denmark; Institució Catalana de Recerca i Estudis Avancats (ICREA), 08010 Barcelona, Spain

## Abstract

The alginate-degrading enzyme *Bo*PL38 of the human gut bacterium *Bacteroides ovatus* CP926 degrades the three polysaccharide structures found in alginate, a major constituent of brown macroalgae with numerous industrial applications. However, the detailed mechanisms of alginate-degrading enzymes remain unclear. Crystal structures of *Bo*PL38 complexes with alginate oligosaccharides, now shed light on the enzyme’s catalytic machinery. QM/MM simulations reveal distinct conformational and reaction pathways, highlighting different transition states for mannuronate and guluronate conversion. C5 proton abstraction at subsite +1 by Y298 and H243 facilitates *syn-* and *anti*-β-elimination reactions, respectively. Substrate recognition relies on R292 distorting the sugar at subsite +1 into a preactivated conformation, while stabilizing the active site tunnel through a salt bridge. Furthermore, NMR spectroscopy found that *Bo*PL38 also catalyze mannuronate to guluronate epimerization in addition to its lyase function, thereby paving the way for future enzymatic alginate modification.

## Introduction

Alginate is a prominent marine polysaccharide, primarily found in cell walls of brown algae^1^. It consists of β-D-mannuronate (M) and its C5 epimer α-L-guluronate (G), linked *via* 1,4-glycosidic bonds, forming homopolyuronate M (polyM) and G (polyG) blocks as well as mixed polyMG blocks^1–3^. This structurally intricate biopolymer has ecological, biomedical, and biotechnological interest due to its uronate composition^4,5^, conferring extraordinary gelation, thickening, and stabilizing abilities. Hundreds of biotechnological applications of alginate are known, ranging from welding rod coating, food industry to drug delivery^5–7^. The wide presence of alginate in marine environments underlines its ecological importance.

Alginate lyases (ALs) depolymerize alginate into alginate oligosaccharides (AOSs) and monosaccharides, which bacteria can assimilate through central metabolic pathways^8,9^, also benefiting their animal and human hosts^10–12^. Understanding the molecular mechanisms and substrate specificity of ALs forming AOSs of defined degrees of polymerization (DP)^12,13^ is pivotal for deciphering the intricacies of alginate metabolism and expanding the potential of the polymer within biotechnological applications.

Based on their amino acid sequence, ALs are currently classified into 15 polysaccharide lyase (PL) families in the carbohydrate-active enzyme database, CAZy (www.cazy.org)^14^. ALs specifically cleave M-M, G-G, M-G, or G-M linkages or combinations thereof^15–17^ (Fig. 1a). More than 10,000 sequences encoding PLs are annotated as prospective ALs, but evidence for enzyme activity exists for less than 200 candidates^14^. Bridging the gap between sequence-based annotation and experimentally verified specificity is necessary for understanding ALs in marine carbon recycling and tailored production of AOSs.

**Fig. 1.**
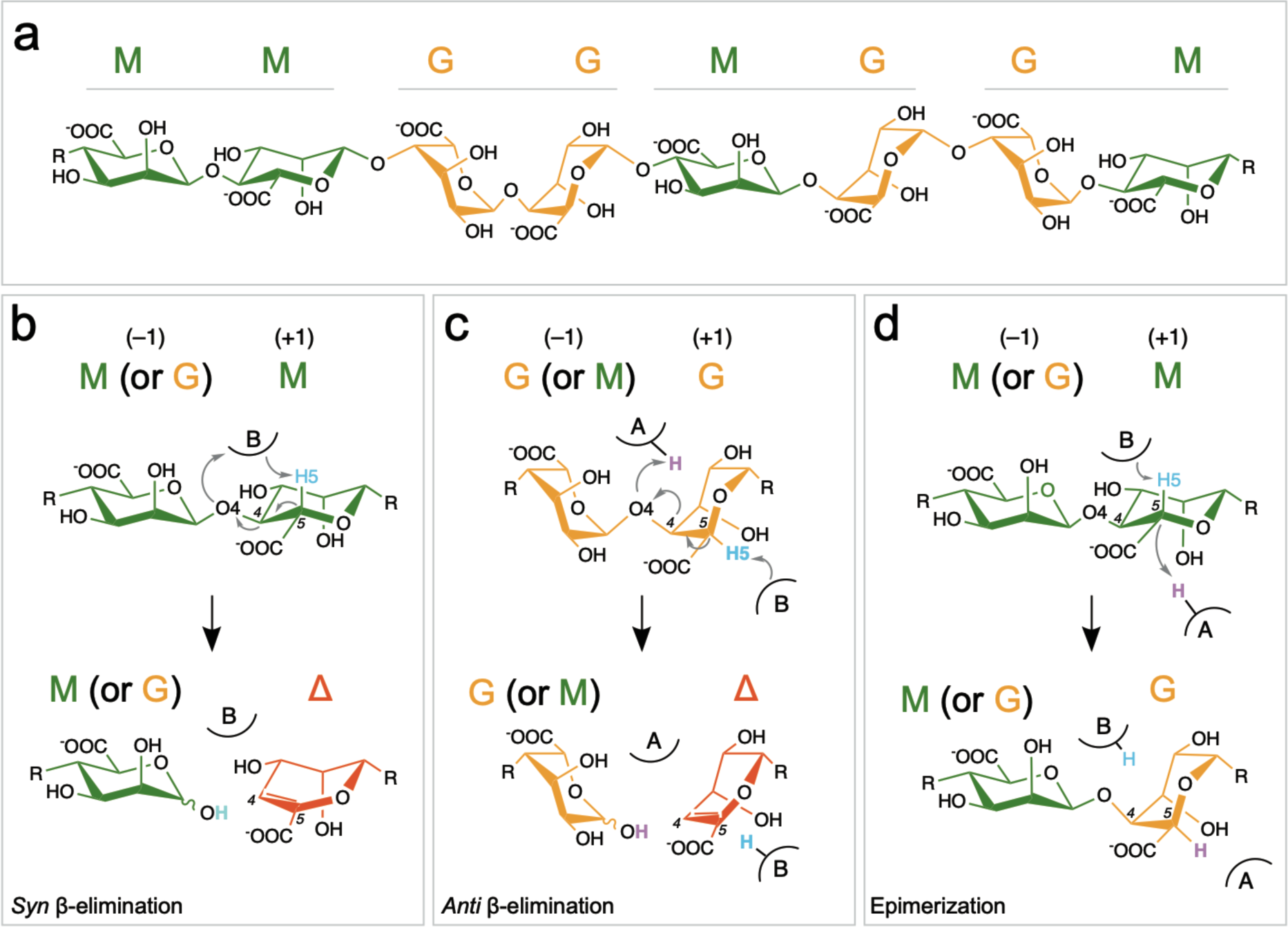
Alginate structure and possible chemical reactions catalyzed by alginate-modifying enzymes. **a** Alginate is made from blocks of mannuronate (M, green), guluronate (G, orange), or blocks of alternating M-G or G-M linkages, here illustrated with dimers representing each block. **b** Cleavage of M-M linkages, such as found in polyM (or G-M linkages of polyMG), is expected to follow a *syn*-β-elimination reaction, where proton abstraction from C5 (H5 in blue) and proton donation to O4 take place on the same side of the substrate, forming a new reducing end M (or G if the initial reactant link is G-M) and a Δ (red) at the non-reducing end. **c** Cleavage of G-G linkages of polyG (or M-G linkages of polyMG) is expected to follow an *anti*-β-elimination reaction, where proton abstraction and donation take place on opposite sides of the substrate. The reaction requires an additional proton (magenta) and results in a new reducing end G (or M if the initial reactant is M-G) and a Δ at the non-reducing end. **d** Similar to the *syn*-β-elimination reaction, the epimerase reaction starts with C5 proton abstraction (H5 in blue). However, instead of a proton being donated to O4, another proton (magenta) is donated to C5 on the opposite side of the substrate, converting M into G. Enzyme subsites –1 and +1 are indicated in panels **b–d**.

The decomposition of alginate by ALs is suggested to occur through a β-elimination mechanism^18^, creating an unsaturated non-reducing end 4-deoxy-L-*erythro*-4-hexenopyranuronate at subsite +1 (denoted by Δ)^19,20^. In this mechanism, a catalytic base (usually a negatively charged tyrosine or a deprotonated histidine) abstracts the H5 proton from the sugar moiety at subsite +1, leading to an unsaturated bond between C4 and C5. The reaction is assisted by proton transfer from a general acid to the glycosidic oxygen (Fig. 1b and c). This two-proton transfer process can take place from the same side (*syn*) or opposing sides (*anti*) of the polymer^21^. In *syn*, the general base abstracts the H5 proton from the uronate moiety at subsite +1 on the same side as the glycosidic oxygen, while in *anti*, they are on the opposite sides^22^. In the *syn* reaction (Fig. 1b), a single residue or two can act as both the general base and acid when M-M and G-M linkages are cleaved because the C5 proton and the departing oxygen are in *syn* relative to each other (Fig. 1b), as recently described for two PL7^23,24^. In contrast, M-G and G-G linkages have the C5 and O4 atoms on opposite sides, thus the *anti-*mechanism is operative (Fig. 1c). The order of reactive events (proton transfers and glycosidic bond cleavage), and the detailed molecular mechanism remains unknown for ALs acting on M-G and G-G links.

ALs can be *endo-*lytic, cleaving alginate chains internally to produce AOS products mixtures of varying DP, or *exo-*lytic, cleaving consistently from the end of the alginate chain, producing monomers (including Δ), or shorter AOSs, such as dimers. ALs may act in either a processive or non-processive manner, which for CAZymes, in general, has previously been linked to fold architecture and the shape of the active site^25^.

Alginate is produced in nature as pure polyM by alginate synthase, and individual M units are converted to G, catalyzed by mannuronate C5 epimerases^26^. As in the lyase-catalyzed β-elimination, the epimerization reaction starts with H5 abstraction^27^. However, instead of cleaving the glycosidic linkage, epimerases change the chirality of the C5 carbon atom of M (Fig. 1d). The alginate epimerases *Av*AlgE7 from *Azotobacter vinelandii*, *Ac*AlgE2, and *Ac*AlgE3 from *Azotobacter chroococcum*^28^, and *Pm*C5A from *Pseudomonas mendocina*^29^ have been shown to display secondary lyase activity^15,27^. However, secondary epimerase activity has not yet been reported in any AL.

Recently, we described the AL *Bo*PL38, belonging to family PL38, from the human gut bacterium *Bacteroides ovatus* strain CP926^30^. *Bo*PL38 is secreted as the primary enzyme degrading alginate into AOSs, enabling *B. ovatus* CP926 to metabolize alginate in its native human gut environment^30^. Unprecedented for ALs, *Bo*PL38 shows remarkable substrate promiscuity, degrading polyM, polyG, and polyMG with similar activity^30^. Understanding the machinery responsible for this diverse substrate preference will expand fundamental insights into the mode of action of ALs.

Here, we combine X-ray crystallography, NMR spectroscopy, molecular dynamics (MD), quantum mechanics/molecular mechanics (QM/MM), and structure-guided mutational analysis to uncover the determinants of substrate promiscuity and the catalytic mechanisms of *Bo*PL38. We reveal the recognition mechanism by which the enzyme exploits the conformational flexibility of the substrate to accommodate different block structures of alginate through distortion of the sugar unit at subsite +1. This facilitates the *syn*- and *anti-*β-elimination reactions to take place at the same enzyme subsite. Moreover, we reveal that the enzyme exhibits secondary epimerase activity, highlighting the extraordinary diversity in the catalytic activity of *Bo*PL38 on AOSs.

## Results

### BoPL38 substrate specificity

Structures of Michaelis-complexes, obtained by X-ray crystallography, of *Bo*PL38 together with hexaguluronate (G_6_), hexamannuronate (M_6_), and hexaoligomeric alternating M-G (*i.e.* (MG)_3_) confirm the location of the active site of the enzyme in the electropositive tunnel of its (⍺/⍺)_7_-barrel fold^30^ (Fig. 2a–b). The complexes were formed at pH 3.5, far from the activity optimum at pH 7.5 of *Bo*PL38 (Extended Data Fig. 1a), with G_4_, M_4_ and (MG)_2_M visible in the electron density of the resulting structures, respectively (Fig. 2c and statistics in Supplementary Table 1).

**Fig. 2.**
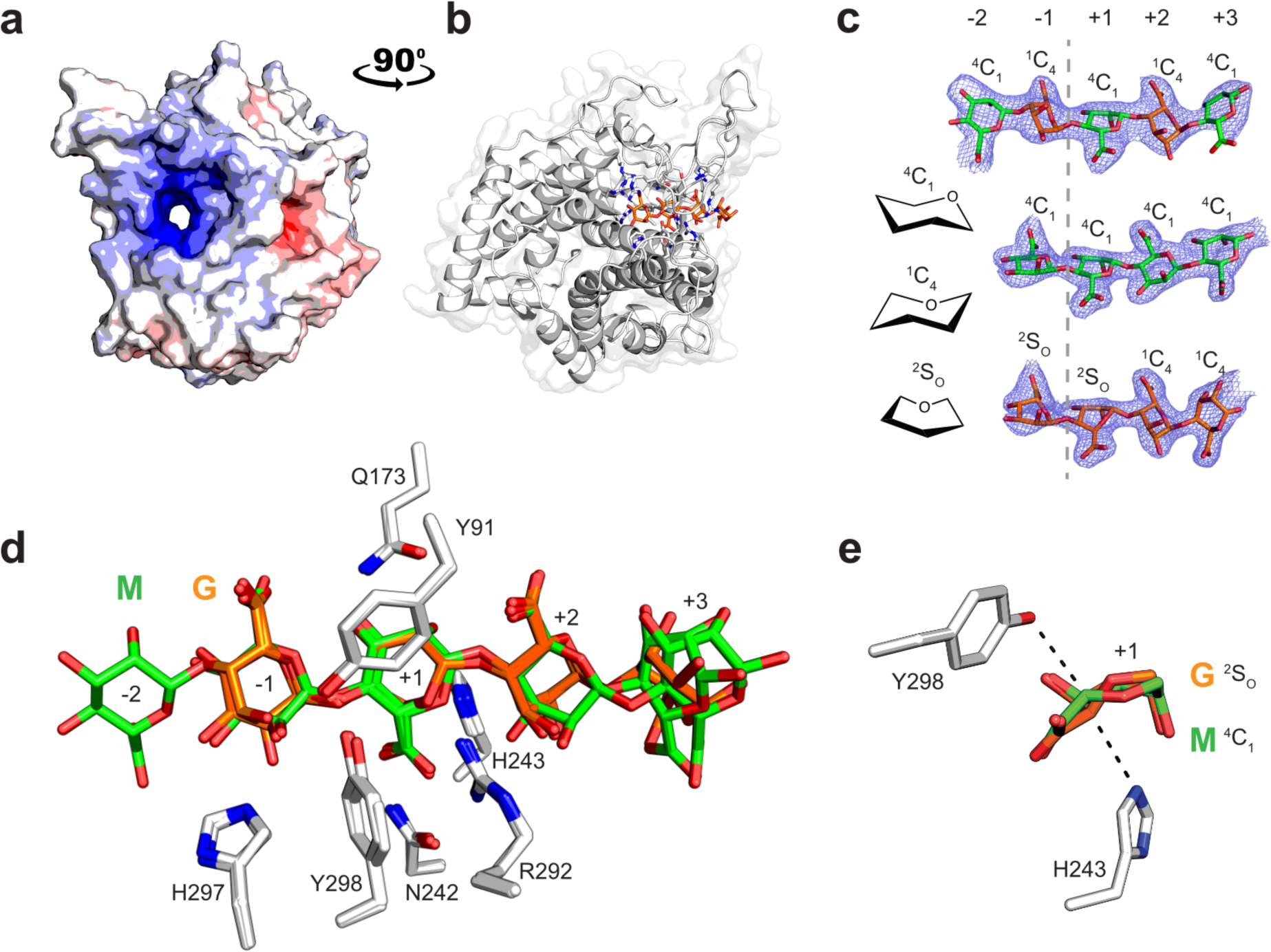
Overall structural features and substrate interactions of *Bo*PL38. **a-b** Surface electrostatics and cartoon representation of *Bo*PL38-G_4_ (substrate not shown in **a**). The two panels are turned 90° relative to each other along the y-axis. **c** Complexes revealed G_4_ (PDB 9FHT), M_4_ (PDB 9FHU), and (MG)_2_M (PDB 9FHV) oligomers binding at the same subsites, and in most cases in their most favorable ^1^C_4_ (G) or ^4^C_1_ (M) conformations. G in subsites –1 and +1 are found in the ^2^S_O_ conformation. Electron density is displayed as 2F_0_-F_C_ in blue mesh at 1.0 σ. **d** Superimposition of the three complex structures, evidencing the near identical orientation of the +1 sugars (see also Extended Data Fig. 1b–c for an extended view). **e** End-on view of subsite +1, indicating the amino acid residues that are likely responsible for proton abstraction from M (Y298, *syn*) or G (H243, *anti*). M residues are shown in green, and G residues in orange.

Most M saccharide units in the enzyme complexes with M_4_ and (MG)_2_M adopt the ^4^C_1_ (canonical chair) conformation, favored in solution^31^. However, a slight distortion towards the ^2^S_O_ (skew-boat) conformation is observed for the M bound in the +1 subsite of M_4_ and (MG)_2_M complexes (Fig. 2c, Supplementary Fig. 1, and Supplementary Table 2). Likewise, the G units in subsites +2 and +3 of the G_4_ complex adopt the ^1^C_4_ (inverted chair) conformation, which is favored in solution, while G units at subsites –1 and +1 are clearly in a ^2^S_O_ distorted conformation (Fig. 2c, Supplementary Fig. 1 and Supplementary Table 2), indicating that the enzyme distorts the conformation of certain sugars upon binding.

The *Bo*PL38 complexes provide clues on the identity of the amino acid residues enabling catalysis of the stereochemically different sugar substrates (Fig. 2d and Extended Data Fig. 1b–c). Structural overlay of the three complexes shows no meaningful differences in conformation of residues closest to subsites –1/+1 (RMSD ≤ 0.157 Å comparing Y91, Q173, N242, H243, R292, H297, and Y298, Fig. 2d). Furthermore, the three complexes and free *Bo*PL38 have similar active sites structures (highest RMSD is 0.26 Å). This corroborates that changes in conformation and orientation occur only on the substrate and not on the enzyme itself.

The conformation of the sugars at subsite +1 (^4^C_1_ for M and ^2^S_O_ for G) result in the C5, and therefore the C5-H5 bond, pointing in opposite directions in M *vs.* G (Fig. 2e). Consequently, the general base abstracting the H5 proton at subsite +1 cannot be the same for G- and M-substrates. In the latter case, Y298, situated above the ring plane of the M sugar, is likely to act as the catalytic base, initiating the *syn*-proton abstraction (Fig. 1b). For G-based substrates, the catalytic base is expected to be H243, located below the ring plane of the G sugar (Fig. 1c), enabling the *anti*-proton abstraction.

To confirm the assignment of Y298 and H243 as catalytic residues, and thereby the site of cleavage, crystals soaked in G_4_ or M_4_ were catalytically activated *via* a 2-min mother-liquor adjustment to pH 8.0. The resulting complex structures are identical to their pH 3.5 counterparts (Fig. 2), except for discontinuous electron density and difference density between subsites –1 and +1 (Extended Data Fig. 1d–e). This substantiates the catalytic roles of Y298 and H243, and the location of the cleavage site between the third and fourth sugar from the reducing end in the G_4_ and M_4_ complexes.

### One active site, two distinct mechanisms

The structures of *Bo*PL38 in complex with G_4_ and M_4_ at pH 8.0 were employed as models for conducting MD simulations of 1.5 µs. The simulations showed that these Michaelis-complexes maintained a catalytically productive configuration (Extended Data Fig. 2a and c). In the G_4_ complex, the H5 atom of the sugar at subsite +1 is poised for proton abstraction, being oriented towards the N_ε_ atom of H243 (Extended Data Fig. 2b). The p*K_a_* of H243 is raised through a strong hydrogen bond between its N_δ_ atom and E236 (Extended Data Fig. 2b and d). In the M_4_ complex, the H5 atom of the sugar at subsite +1 is ready for proton abstraction by the O_ƞ_ of Y298 (Extended Data Fig. 1d). This corroborates H243 and Y298 serving as the catalytic bases for the *anti*- and *syn*-β-elimination reactions with polyG and polyM, respectively.

A representative snapshot from the classical MD simulations of each *Bo*PL38 complex was chosen to initiate QM/MM MD simulations of the reaction mechanisms. A large QM region, including the substrate (at subsites –1, +1, and part of subsite +2) and the sidechains of R292, E236, H243, and Y298, was used for each system (Extended Data Fig. 3a–b). QM/MM simulations, using the OPES enhanced sampling method^32^, successfully drove the system from reactants (*Bo*PL38 in complex with either M_4_ or G_4_) to products (*Bo*PL38 in complex with M and Δ-MM or G and Δ-GG). The QM/MM OPES simulations were guided by a path collective variable (PathCV)^33,34^ accounting for all anticipated chemical movements along the reaction coordinate, including proton transfer events and covalent bond changes (Extended Data Fig. 3c–f).

The free energy profile for G_4_ conversion (Fig. 3) shows two main minima (reactants and products, with the latter being lower in energy) separated by a single transition state (TS), indicating an exothermic and concerted (*i.e.* one step) reaction. The computed^35^ and experimentally estimated^30^ free energy barriers correlate well (18.5 kcal mol⁻¹ *vs. k_cat_* 9.1 ± 0.11 s⁻¹, 16.8 kcal mol⁻¹), indicating a feasible *anti*-β-elimination reaction for *Bo*PL38 (Fig. 1c).

**Fig. 3.**
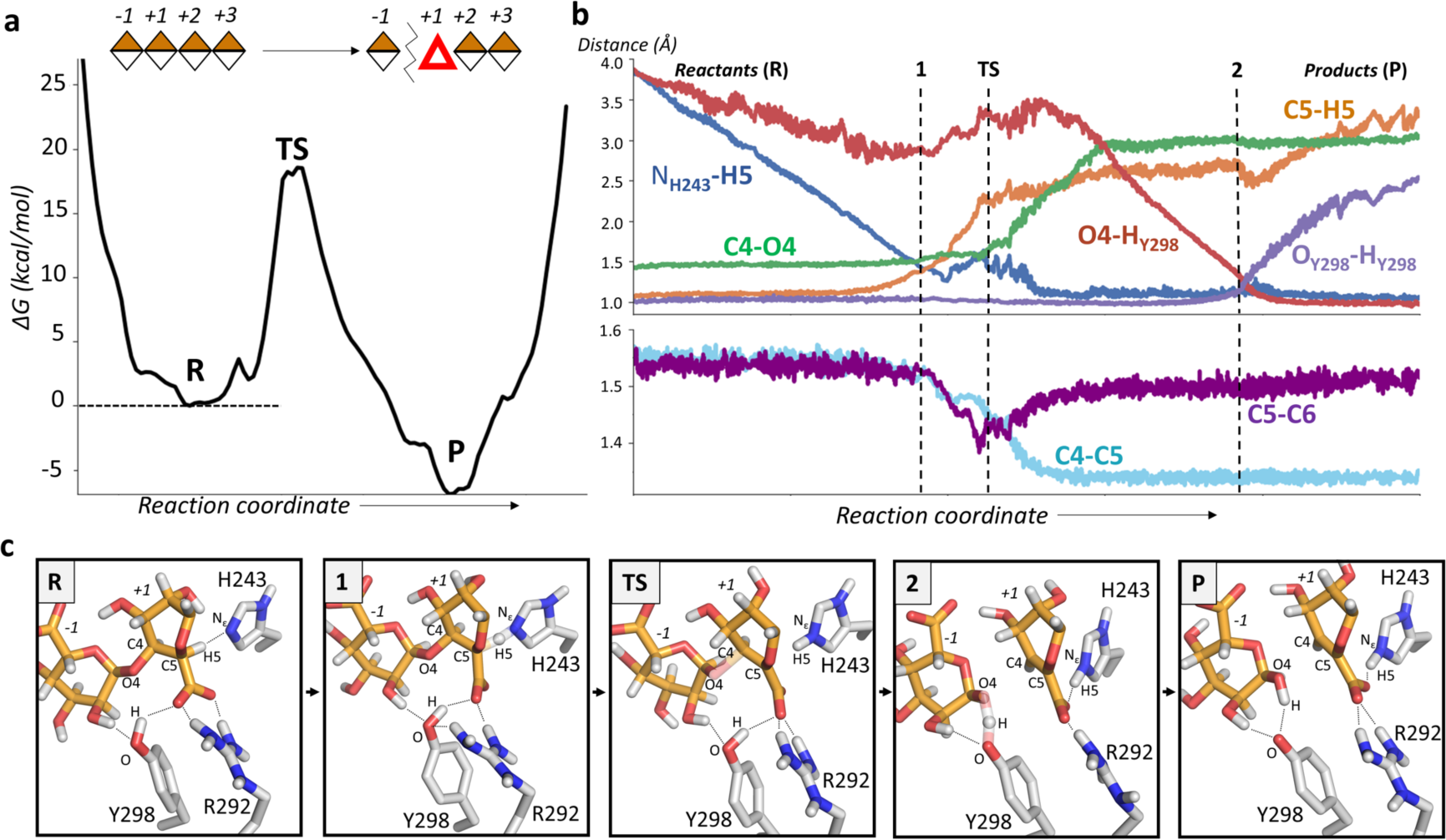
A*n*ti-β-elimination reaction mechanism of *Bo*PL38 on G_4_. **a** Free energy profile (kcal mol⁻¹) of the reaction mechanism along the reaction coordinate. **b** Evolution of the main catalytic distances through the reaction coordinate. **c** Representative structures of the active site along the reaction coordinate. Only the –1 and +1 sugars of the G_4_ substrate are shown. Non-polar hydrogen atoms have been omitted for clarity, except the H5 atom at subsite +1. Hydrogen bonds are represented with black dotted lines, while transparent bonds represent partially broken/formed. After the reaction, O4 becomes O1 of the G unit bound at subsite –1 (not renamed in the picture). The reaction shown in panel **c** is also presented as Supplementary Video 1.

Analysis of the molecular changes along the reaction coordinate for conversion of G_4_ shows that the initial ^2^S_O_ conformation of the sugar at subsite +1 enables a close contact between its H5 atom and H243 (Fig. 3cR). Subsequently, H243 abstracts the H5 proton (Fig. 3c1), and the C5 atom acquires a negative charge, which delocalizes over C4. This causes the C5-C4 bond to shrink (forming a partial double bond) and the glycosidic bond to elongate. At the TS, the glycosidic bond is broken and a double bond between C4 and C5 begins to form. Subsequently, residue Y298 serves as the catalytic acid, donating its proton to the new reducing end (Fig. 3c2), leading to the formation of the reaction products G and Δ-GG (Fig. 3cP). At this point, the Δ sugar at subsite +1 adopts a B_3,0_ conformation, most likely due to hydrogen bond interactions with enzyme residues Q173, H243, and R292 (Fig. 3c and Extended Data Fig. 3g). This contrasts with the ^2^E conformation, recently reported for PL7^23^, evidencing that enzyme-specific interactions with the substrate lead to different product conformations.

The simulations of *Bo*PL38 reacting with M_4_ confirm the expected *syn*-β-elimination mechanism. As with G_4_, the reaction is concerted and exergonic (Fig. 4a), with a similar free energy barrier (≈ 20 kcal mol⁻¹) agreeing with experimental values^30,35^ (*k_cat_* of 0.9 ± 0.02 s^–1^, 18.2 kcal mol⁻¹). Analysis of the atomic motion along the minimum free energy pathway shows R292 distorting the bound sugar, bringing the H5 proton close to Y298 for proton abstraction (Fig. 4c1). Dissimilar to the G_4_ reaction, the glycosidic linkage is still maintained at the TS, and the sugar at subsite +1 shows a higher carbanion character (negative charge accumulated at C5). After the TS, the glycosidic bond breaks and Y298 transfers the proton to the new reducing end (Fig. 4c2 and 4cP, and Extended Data Fig. 3h), leading to the reaction products. Similar to the G_4_ reaction, substrate-enzyme interactions restrict the conformations of M (and subsequently Δ) at subsite +1, limiting it to ^2^S_O_ and B_3,O_ conformations throughout the catalytic process.

**Fig. 4.**
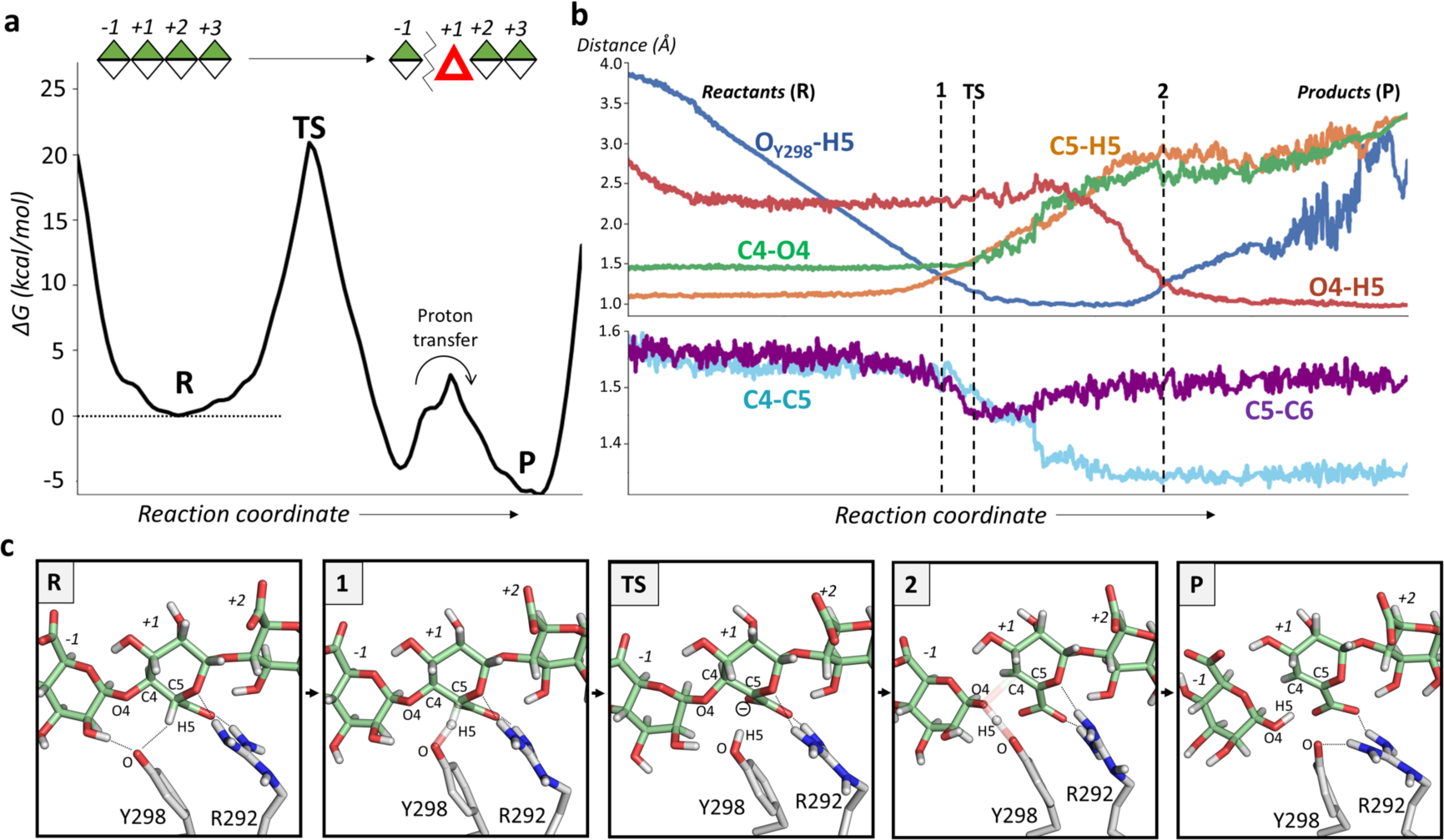
S*y*n-β-elimination reaction mechanism of *Bo*PL38 on M_4_. **a** Reaction free energy profile. Energies are reported in kcal mol⁻¹. The arrow indicates the proton transfer from Y298 to the sugar at the –1 subsite. **b** Evolution of the main catalytic distances along the reaction coordinate. **c** Representative active site structures along the reaction coordinate. Only the –1 and +1 sugars of the M_4_ substrate are shown. Non-polar hydrogen atoms have been omitted for clarity, except the H5 atom at subsite +1. Hydrogen bonds are represented with black dotted lines, while transparent bonds are partially broken/formed. After the reaction, O4 becomes O1 of the M unit at subsite –1 (not renamed in the picture). Panel **c** is also presented as Supplementary Video 2.

### NMR reveals secondary epimerization activity

Time-resolved NMR spectroscopy is used to confirm the identity of the reaction products, verify the preference for G-M *vs.* M-G linkages, and determine the mode of action (*exo-*lytic *vs*. *endo-*lytic) as *Bo*PL38 reacts on polyG, polyM, and polyMG. A complete assignment of NMR signals for the two reference compounds, Δ-MM and Δ-GG, under the same conditions as the *Bo*PL38 reactions (Supplementary Fig. 2, Supplementary Tables 3–4) is used to assign the NMR signals for *Bo*PL38 reactions with various substrates (Fig. 5, Supplementary Fig. 3–5, Supplementary Tables 5–6).

**Fig. 5.**
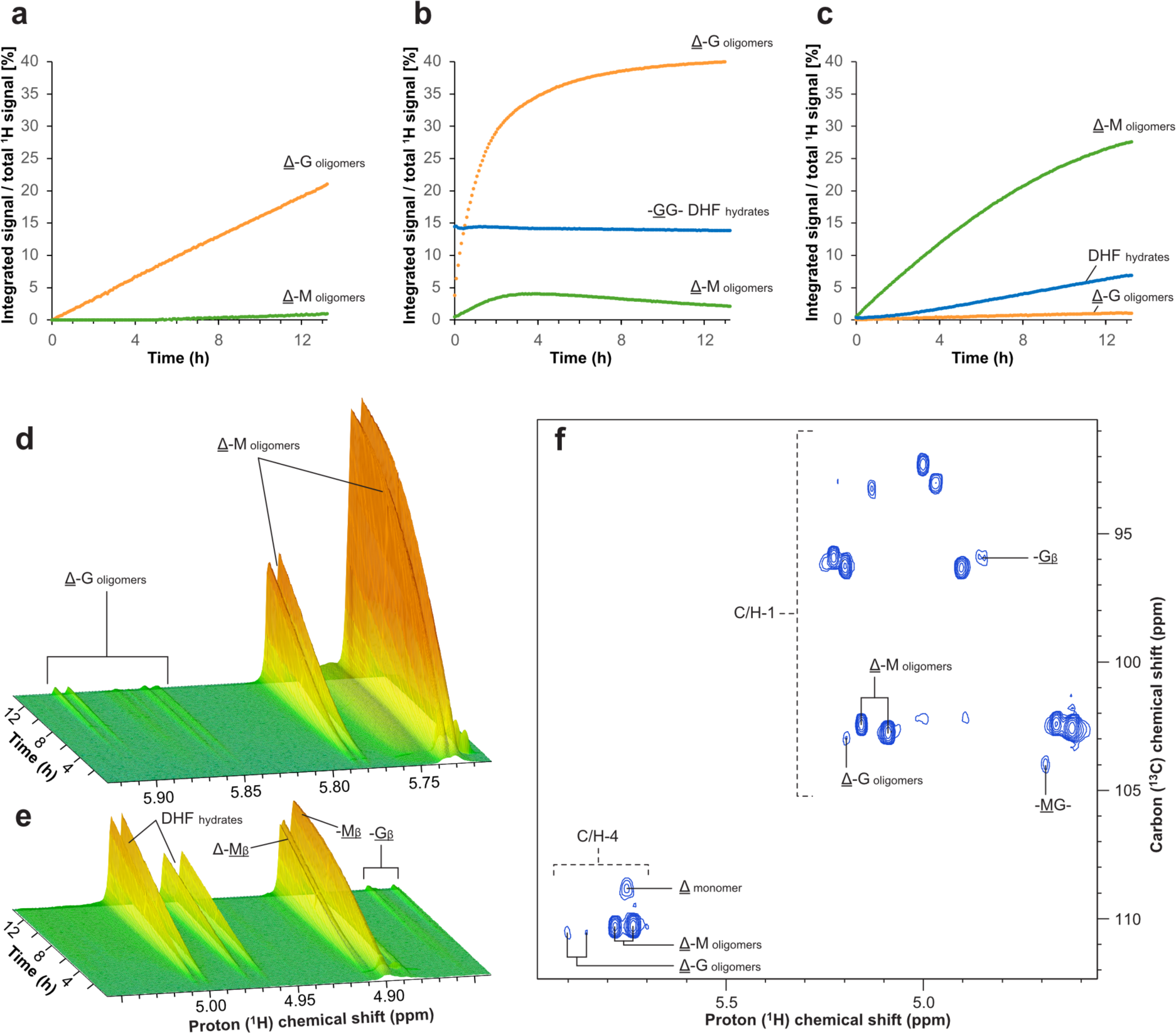
Time-resolved NMR analysis for reactions with *Bo*PL38 and polyMG, polyG, and polyM. Regions for lyase reaction products were integrated and plotted through time (relative to the total ^1^H signal at the end of the reaction) for *Bo*PL38 reacting with polyMG, polyG, and polyM. **a** The reaction with polyMG reveals the formation of mainly Δ-G dimers and oligomers with little product of Δ-M character formed. **b** The reaction with polyG produces mainly Δ-G dimers and oligomers (Δ-M products were also observed due to residual M in the initial polyG substrate, F_G_>0.95). Monomer products are also observed as DHF. Due to the overlap of the DHF signals with the substrate -GG- signal, the indicated percentage of DHF formation is not reliable. The HSQC recorded after the reaction clearly showed that DHF had been formed. **c** Time-resolved analysis of *Bo*PL38 with polyM reveals both monomer DHF hydrates and Δ-M and Δ-G oligomers resulting from lyase activity. **d, e** The time-resolved NMR spectra of the *Bo*PL38 reaction on polyM show Δ-M and Δ-G oligomers formed by the lyase activity and G-reducing ends associated with epimerization activity for the two panels, respectively. **f** HSQC spectrum of the reaction on polyM shows correlations between directly bonded H-C, lyase products (Δ, Δ-M oligomers and Δ-G oligomers), and epimerization products (G_β_-reducing ends and internal M-G signals). The complete assignment of NMR spectra is shown in Supplementary Fig. 3–5.

*Bo*PL38 reacting with polyMG (with a fraction of M (F_M_) = 0.5, Supplementary Fig. 6a) produces 4,5-unsaturated products, confirming lyase activity. The products are almost exclusively Δ-G oligomers and Δ-G dimers with no detectable monomers (Fig. 5a and Supplementary Fig. 3), indicating a strong G-M bond preference, with minor signals corresponding to cleaved M-G bonds. Reaction with polyG (F_M_ < 0.15, Supplementary Fig. 6b) shows the main lyase products are Δ-G dimers but also monomers of Δ (Fig. 5b and Supplementary Fig. 4).

When degrading pure polyM (F_M_ = 1, Supplementary Fig. 6c), Δ-M dimers are produced along with Δ monomers, confirming lyase activity on M-M bonds (Fig. 5c and in more detail in Supplementary Fig. 5). However, unexpectedly, weak signals of Δ-G products, internal M-G bonds, and G_β_-reducing ends are also observed (Fig. 5d–f). Since the substrate consists of pure polyM, the occurrence of G-residues can only be explained by sugar epimerization followed by lyase activity.

The mode of action was further analyzed with HPAEC-PAD, which reveals a broad distribution of products from polyG, polyM, and polyMG substrates (Extended Data Fig. 4a-c). Regardless of the substrate, *Bo*PL38 operates preferably in *endo* mode, producing primarily dimers and, to a small extent, monomers of Δ, M or G, corroborating both crystal structures (Fig. 2) and computational models of the reaction (Fig. 3–4).

The hydrolytic ring-opening of Δ to 4-deoxy-L-*erythro*-5-hexulosuronate (DEH) and rearrangement into stereoisomers of 4-deoxy-D-manno-hexulofuranosidonate (DHF) is relevant for bacterial carbohydrate uptake^20,36^. Both DEH and DHF hydrates are noted as reaction products (Fig. 5, Supplementary Fig. 3–5 and Extended Data Fig. 4a–c). Soaking *Bo*PL38 crystals with unsaturated tetrasaccharides (Δ-MMM and Δ-GGG) resulted in bound Δ-MM and Δ-GG oligomers with additional electron density fitting unlinked Δ or DHF (Supplementary Fig. 7), which correlates well with NMR observations.

### Site-directed mutagenesis

*Bo*PL38 variants of nine active site residues (Table 1) were tested for lyase activity on polyM and polyG *via* the A_235 nm_ spectrophotometric assay of the formation of unsaturated chain ends (Extended Data Fig. 4d, Supplementary Fig. 8), and in addition for lyase and epimerase activity using NMR spectrometry (Extended Data Fig. 4e, Supplementary Fig. 9, and Supplementary Tables 8–9). The results for each *Bo*PL38 variant, including the predicted functional role of the original residue, are summarized in Table 1.

**Table 1.**
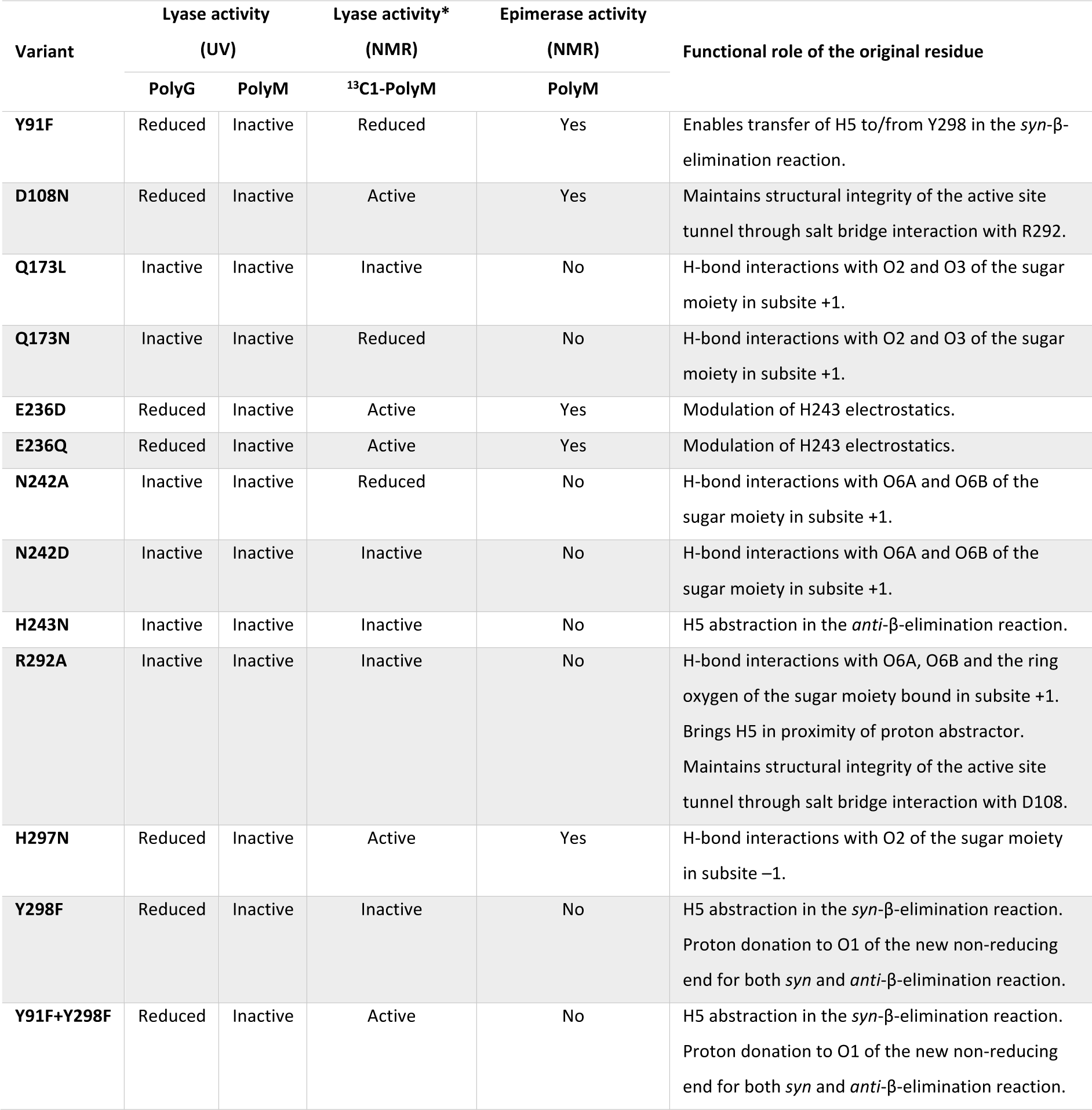
Summary of lyase and epimerase activity of *Bo*PL38 variants. The variants are listed along with their phenotype with regard to lyase and epimerase catalysis and the hypothesis for the functional role of the original residue. _*_ Active means >0.5% WT activity. Reduced means <0.5% WT activity. Inactive means no observed signal.

*Y298 variants:* Y298F did not exhibit polyM activity, as expected when the general base for the *syn*-reaction is lacking. the crystal structure of *Bo*PL38 Y298F in complex with M_4_ (Supplementary Fig. 10a), indicates that Y298 is not essential for substrate binding.

*H297 variants:* Inspection of the crystal and computed structures led to the hypothesis that H297 is involved in the regulation of Y298 protonation. The H297N mutation reduced activity to 12.5% lyase and 0.19% epimerase activity, compared to the total activity of the WT at 96% lyase and 4% epimerase activity, respectively (Extended Data Fig. 4e). In the complex structures, H297 forms an H-bond to the C2-OH of the sugar bound in subsite –1. H297N is unable to form this interaction and loses the ability to bind the substrate, as demonstrated *in crystallo* (Supplementary Fig. 10b). This indicates that H297 is important for the formation of the Michaelis-complex, rather than having a regulatory role for protonation.

*Y91 variants:* The Michaelis-complex structures indicate that the interaction between Y91 and Y298 could influence the protonation state of Y298 (Fig. 2d–e and Extended Data Fig. 1d–e). The Y91F mutation greatly reduced polyM activity, suggesting Y91 influences protonation of Y298, the catalytic base of the *syn*-reaction. Notably, the Y91F+Y298F variant retained ∼10% lyase and <1% epimerase activity, respectively (Extended Data Fig. 4e), and, like the Y298F variant, it binds substrate *in crystallo* (Supplementary Fig. 10c). Since the catalytic base of the *syn*-elimination is missing, the Y91F+Y298F variant seems to employ some rescue mechanism, only operative if both mutations are in play. Similarly, residual activity on polyG of the Y298F and Y91F+Y298F variants alludes to a partial rescue *anti*-reaction in the absence of the catalytic acid Y298.

*H243 variants:* Mutation of H243 was expected to only affect the enzyme activity on polyG (*anti*-elimination) since this residue plays no role in the *syn*-β-elimination mechanism on polyM (Fig. 3). However, the H243N variant turned out to be inactive on both polyM and polyG (Extended Data Figs. 4d–e and Supplementary Figs. 8–9). These results were rationalized by MD simulations showing that H243N disrupts the network of hydrogen bond interactions at the active site in both cases, causing structural unfolding (Supplementary Fig. 11a) leading to enzyme inactivation.

*E236 variants:* E236 was found to interact with H243 in the MD simulations (Extended Data Fig. 2), suggesting that this residue increases the p*K_a_* of H243, thereby making it relevant for the *anti*-reaction. However, the E236D/Q variants showed greatly reduced polyG and polyM activity (Table 1 and Extended Data Fig. 4e). The MD simulations show that the E236D/Q variant is structurally unstable, which may explain its reduced activity on both substrates (Supplementary Fig. 11b–c).

*N242 variants:* N242 interacts with the uronate carboxylate group at subsite +1 (Fig. 2d). The N242D mutation would make the active site of *Bo*PL38 more similar to that of the AlgE7 epimerase^27^, potentially increasing epimerase activity. However, both epimerase and lyase activities were lost in the N242D/A variants. MD simulations of N242D revealed a significant conformational shift in the active site tunnel (Supplementary Fig. 11d), preventing substrate binding, rationalizing the lack of activity. Similarly, the Q173N/L mutations abolished both lyase and epimerase activities. N242A and Q173N variants have destabilized loop conformations as shown by MD simulations and crystal structures, likely preventing substrate binding (Supplementary Fig. 10d–e and 12).

*R292 variants:* The simulations suggest that R292 is essential for bringing H5 close to Y298 for M-M bond cleavage. Additionally, R292 forms a salt bridge with D108, connecting two loops that create the ‘roof’ of the active site tunnel. MD simulations of *Bo*PL38 WT, D108N, and R292A revealed that disrupting this salt bridge opens the active site and separates the catalytic residues (Supplementary Fig. 13), a point not evident from thermal stability analysis (Supplementary Fig. 14). The R292–D108 interaction is crucial for the structural integrity of the active site tunnel. Experimental data for *Bo*PL38 D108N in the presence of M_4_ matched the MD simulation structure (RMSD = 1.5 Å), confirming its inability to bind the AOS (Supplementary Fig. 15). These results correlate well with the observed complete and partial enzyme inactivation of R292A and D108N, respectively.

## Discussion

Understanding the mechanistic details of enzymes that degrade marine polysaccharides is crucial for optimizing marine carbon recycling and the efficient utilization of macroalgae. This study identifies the molecular determinants underlying the substrate promiscuity and elucidates the catalytic mechanisms of *Bo*PL38, a newly characterized alginate-degrading enzyme^30^. Using a combination of experimental and theoretical techniques, the roles of all active site residues are dissected, and the different mechanisms adopted by *Bo*PL38 to cleave glycosidic linkages of M-M, G-G, and mixed G-M blocks of alginate are uncovered. X-ray crystallography and MD simulations revealed a tunnel-shaped active site formed by two loops connected by a salt bridge between R292 and D108, essential for maintaining the tunnel structure and enzyme activity. Mutations of active site residues not only reduce enzyme-substrate interactions but also compromise the integrity of the catalytic tunnel, highlighting a finely tuned interaction network within the active site.

*Bo*PL38 accommodates polyM and polyG substrates in its tunnel-shaped active site, with most sugar units adopting relaxed chair conformations (^4^C_1_ for M and ^1^C_4_ for G). However, the sugar units at subsites –1 and +1 in the G_4_ complex, and to a lesser extent, at subsite +1 in the M_4_ complex, adopt a distorted ^2^S_O_ conformation. This directs the C5-H5 bond towards the general base catalyst for proton abstraction (H243 for polyG and Y298 for polyM), promoting catalysis. The sugar at subsite +1 interacts with active site residues Q173, N242, and R292, imposing conformational control. Mutations disrupting these interactions abolish lyase activity for both substrates. This leads to the hypothesis that *Bo*PL38 has evolved to accommodate substrates in a preactivated ^2^S_O_ conformation promoting catalysis. This is analogous to the conformational control exerted by glycoside hydrolases (GHs)^37^, where the enzyme binds substrates in a catalytically favorable distorted conformation, though at subsite –1 rather than subsite +1 as in PLs^14,21^.

Following conformational preactivation, the lyase reaction on polyM requires Y298 to be deprotonated to abstract the H5 proton of the M saccharide at subsite +1, while the protonation state of H243 is not relevant for this process (Supplementary Fig. 16a). In contrast, for polyG conversion, Y298 must be protonated, and H243 deprotonated, due to their specific roles acting on the opposite C5-H5 orientation of the G sugar at subsite +1 compared to M, its C5 epimer (Supplementary Fig. 16b). These intricate substrate-specific protonation requirements suggest that the unliganded enzyme exists in distinct populations, each with varying protonation states of Y298 and H243. The substrate selectively binds to the enzyme population in the appropriate state that is optimal for catalysis.

QM/MM simulations confirm that *Bo*PL38 follows a β-elimination mechanism for cleaving glycosidic linkages in both polyM and polyG, with a *syn*-type reaction for polyM and *anti*-type for polyG. The slightly lower activation energy for polyG (18.5 kcal/mol) compared to polyM (20.9 kcal/mol) aligns with experimentally measured reaction rates^30^ (9.1 ± 0.11 s⁻¹ for polyG and 0.9 ± 0.02 s⁻¹ for polyM). Both reactions are concerted but asynchronous, with proton abstraction preceding bond cleavage. At the TS of the reaction on polyG, the glycosidic bond is partially broken, while it remains intact for polyM that exhibits a higher carbanion-like character. A similar TS was recently found for a PL7 family member acting on polyM^23^, suggesting a carbanion-like uronate configuration in polyM-active ALs in general. These differences are probably due to the distinct intrinsic structures of the two polymers. PolyG is highly constrained in the *Bo*PL38 active site, and its structure deviates from its natural rigid structure^38^ (Supplementary Fig. 1a), while polyM remains unaffected due to its inherent extended structure^38^ (Supplementary Fig. 1d, Supplementary Table 2). Consequently, the glycosidic linkage of polyM is less impacted by substrate binding than that of polyG.

It is intriguing to examine the changes in protonation states of the catalytic residues during the reaction, as these changes can reveal the enzyme’s operational mode across distinct reaction cycles. During the reaction on polyG, H243 becomes protonated, thus, proton transfer is needed to restore the catalytic residues for the next catalytic cycle. This indicates that substrate dissociation must allow the active site to recover the appropriate protonation states before substrate rebinding. The orientation of Y298 and H243 toward each other suggests that a water-mediated proton shuttling mechanism is involved (Supplementary Fig. 17). This may explain why H243 mutations, thought irrelevant to the polyM reaction, led to a loss of all enzyme activities.

These findings indicate that *Bo*PL38 acts non-processively on polyG, aligning with the product distribution observed in HPAEC-PAD (Extended Data Fig. 4a). Although one might expect processivity on polyM, given that Y298 remains deprotonated between cycles, the variable distribution in product DP suggests otherwise (Extended Data Fig. 4b). Therefore, *Bo*PL38 must be non-processive on both substrates.

*Bo*PL38 is the primary alginate-degrading enzyme secreted by *B. ovatus* CP926^30^. It works synergistically with the G-specific *Bo*PL6 and M-specific *Bo*PL17, both encoded in the same polysaccharide utilization locus as *Bo*PL38^30^. *Bo*PL38 rapidly generates DP5–7 AOSs, and yields DP2 products over time *in vitro*, as shown by NMR (Supplementary Fig. 3–5). The bacterium transports the AOS products to the periplasm for degradation by *Bo*PL6 and *Bo*PL17 to Δ monomers, subsequently transported into the cytoplasm for metabolism^30^. The broad substrate preference of *Bo*PL38 likely evolved to tackle the scarcity of alginates and other extracellular ALs in the intestinal environment.

While dual epimerase and lyase activity has been shown in alginate epimerases^15,27^, until now, no PL family has been reported to have epimerase activity until now. Alginate epimerase activity is well-documented in parallel β-helix fold proteins like AlgG, AlgE, and PsmE (EC 5.1.3.37)^39^, with processive action shown for *Av*AlgE7, *Ac*AlyE2, and *Ac*AlyE3^27,39^. In contrast, *Bo*PL38 exhibits an (α/α)_7_-barrel, demonstrating that epimerase activity is not restricted to the anti-parallel β-helix. Thus, *Bo*PL38 bridges lyase and epimerase functions, introducing the dual functionality to a distinct structural fold.

The epimerization side activity of *Bo*PL38 is consistent with lyase activity measurements on polyM by the A_235 nm_ assay. The initial “lag phase”, observed previously^30^ (Supplementary Fig. 8), suggests complex kinetics involving a multistep process where epimerization precedes lyase reaction. The balance between a high *k_cat_* of the polyG reaction and a low *K_M_* for polyM likely regulates lyase and epimerase activities. If M was converted to G and immediately cleaved, the epimerization event would remain undetected, as G would convert to Δ, masking any G signals. Thus, product dissociation must occur before the next catalytic cycle.

This discovery of *Bo*PL38’s diverse activities suggests that other lyase families may also harbor catalytic machinery similarly enabling multiple functions. For instance, members of M-specific PL families without proven lyase activity might possess epimerase activity. The selective pressure for epimerase activity in lyases remains unclear and may simply be a consequence of having lyase activity on both G and M sugars. For dual activity, lyases must have a low K_M_ and an amino acid arrangement capable of abstracting H5 from one side of the sugar ring and donating it to the other side. This arrangement, observed in the +1 subsites of *Bo*PL38 and epimerase structures, involves Y298 and H243 in *Bo*PL38 and Y149 and H154 in *Av*AlgE7^27^. Additionally, the carbanion-like TS in the *syn*-elimination reaction on polyM offers a plausible chemical species for M-to-G conversion during epimerization. This could explain the directionality of the epimerization reaction (M to G and not G to M) and its occurrence in M-acting ALs.

The present work renews the knowledge on enzymatic alginate manipulation by dissecting the mechanistic details of the AL *Bo*PL38, its substrate interactions, and products from lyase and epimerase reactions. The amino acid arrangement at the active site in *Bo*PL38 exemplifies a unique structural constellation fulfilling the requirement for dual lyase and epimerase functions exerted in a multispecific AL. This motivates future exploration of the full potential of *Bo*PL38, and the PL38 family in general, both to place this in a biological context, to manipulate the lyase/epimerase activity ratio and to rationally engineer for desirable preference.

## Acknowledgment

This work is supported by Independent Research Fund Denmark | Natural Sciences via the DSMADE project (DFF170746) (TT, MER, MM, and BS), Novo Nordisk Foundation (NNFOC0027616) (MM and BS), the Technical University of Denmark (TT, MER, MM, CW, and BS), Norwegian Research Council (RCN) grant 315385 (AlgModE) (ABP, LJK, FLA), and grant 226244 (Norwegian NMR Platform) (ABP, LJK, FLA), and the Spanish Ministry of Science, Innovation and Universities (MICIU/AEI/10.13039/501100011033 and FEDER, UE, grant PID2023-147939NB-I00, to CR), the Spanish Structures of Excellence María de Maeztu (MICIU/AEI/10.13039/501100011033 grant CEX2021-001202-M, to CR), the Agency for Management of University and Research Grants of Catalonia (AGAUR, grant 2021-SGR-00680, to CR) and the European Research Council (grant ERC-2020-SyG-951231 “Carbocentre”, to CR). JPR was supported by MICINN (FPI fellowship PRE2021-097898). We thank the Danish Agency for Science, Technology, and Innovation for funding the instrument center DanScatt, which supported our usage of beam time at the synchrotrons. MAX IV Laboratory is acknowledged for time on beamline BioMAX under Proposal 20200120. Research conducted at MAX IV, a Swedish national user facility, is supported by the Swedish Research Council under contract 2018-07152, the Swedish Governmental Agency for Innovation Systems under contract 2018-04969, and Formas under contract 2019-02496. We also acknowledge the European Synchrotron Radiation Facility for the provision of beam time on ID30A3, ID30B, ID23-1, and ID23-2 and would like to thank Andrew McCarthy, David Flot, Gianluca Santoni, Romain Talon, and Antoine Royant for their assistance. The Mosquito crystallization robot was granted by the Carlsberg Foundation and used for obtaining some of the protein crystals. We thank Olav A. Aarstad, Karina Jansen, Melisa Hamzic, and Mivan N. H. Saleh for laboratory assistance.

## Data availability

All crystal structures have been deposited to the Protein Data Bank (www.rcsb.org). Simulation data can be found in the Zenodo repository (DOI: 10.5281/zenodo.12793724). These will be released upon peer-review publication.

## Extended Data

**Extended Data Fig. 1.**
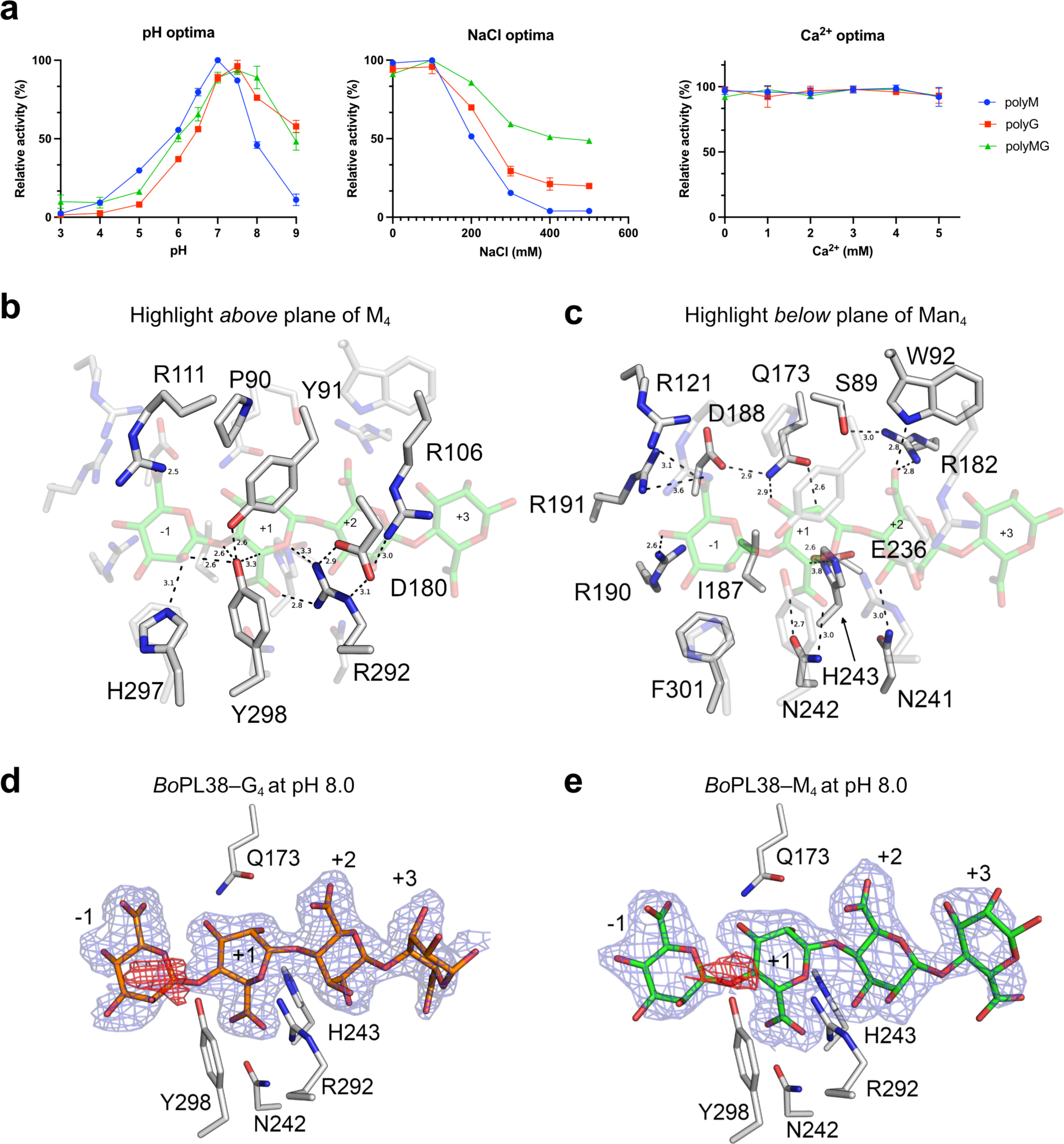
Reaction condition optima and active site residues of *Bo*PL38. **a** pH, NaCl and Ca^2+^ optimal for *Bo*PL38 reacting on polyM, polyG or polyMG. **b** The amino acid residues located above and **c** below or in plane with the bound sugar are highlighted in *Bo*PL38-M_4_ (PDB 9FHU). The panels are in the same orientation. Distances are shown in Å. **d** Discontinuous electron density (blue mesh) and negative difference density (red mesh) indicates cleaved substrate in the *Bo*PL38 structures solved at pH 8.0 of G_4_ (PDB 9FHW) and **e** M_4_ (PDB 9FHX) complexes. Electron density is displayed as 2F_O_-F_C_ in blue mesh at 1.0 σ. Difference density is shown as F_O_-F_C_ in green and red mesh at ±3.0 σ. M residues are shown in green, G residues in orange.

**Extended Data Fig. 2.**
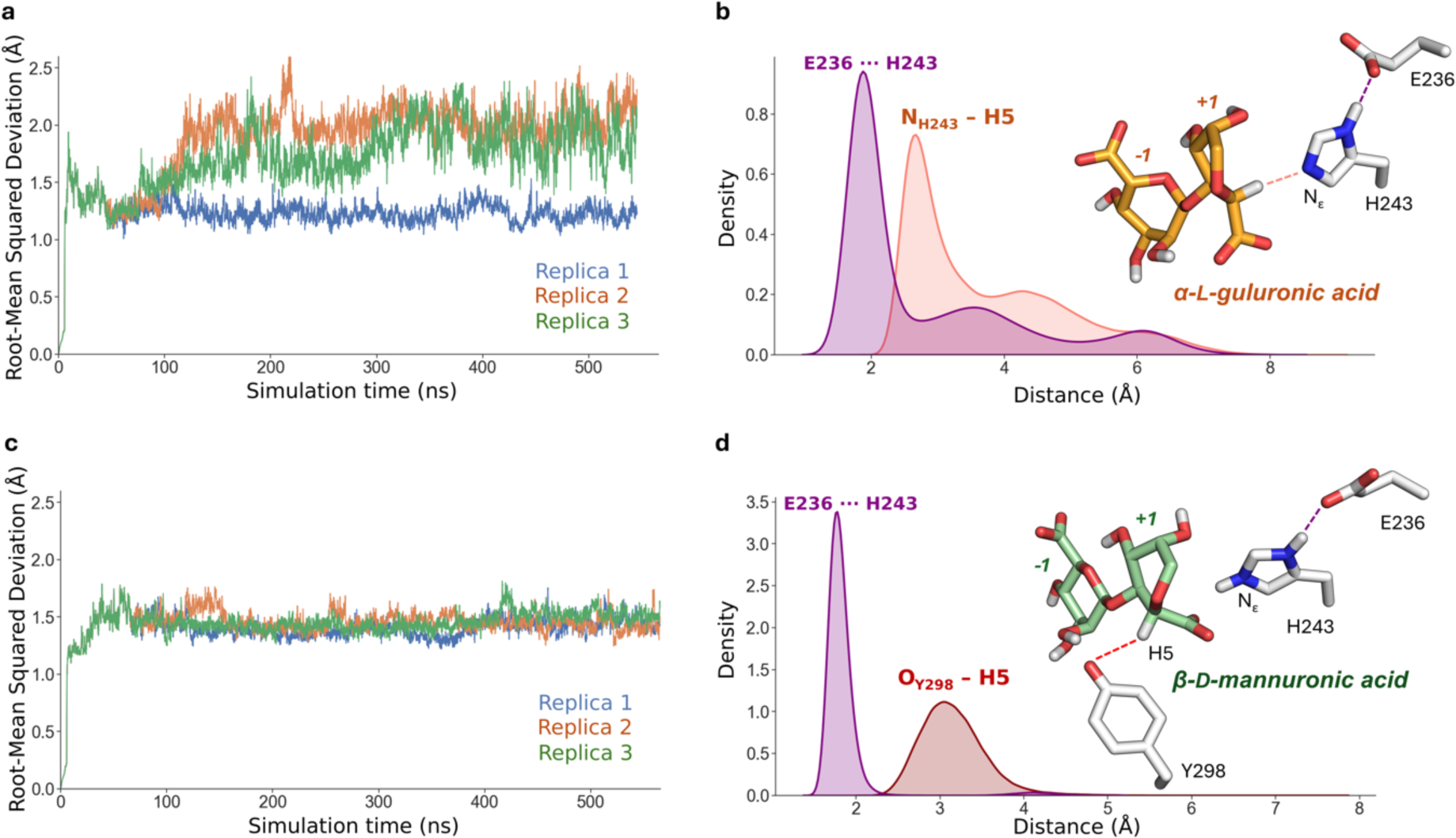
Analysis of the stability of classical simulations of *Bo*PL38 in complex with G_4_ and M_4_. **a** RMSD of the cMD simulations on the G_4_ complex. Different colors were used for the different replicas. **b** Distribution of distances of the nucleophilic attack (N_ε_···H5) and during the cMD simulations with G_4_. The simulation elucidates the interaction E236···H242, increasing the p*K_a_* of H242. **c** RMSD of the cMD simulations on the M_4_ complex. Different colors were used for the different replicas. **d** Distribution of distances of the nucleophilic attack (O_Y298_···H5) during the cMD simulations. Also here, the simulation elucidates the interaction E236···H242, increasing the p*K_a_* of H242.

**Extended Data Fig. 3.**
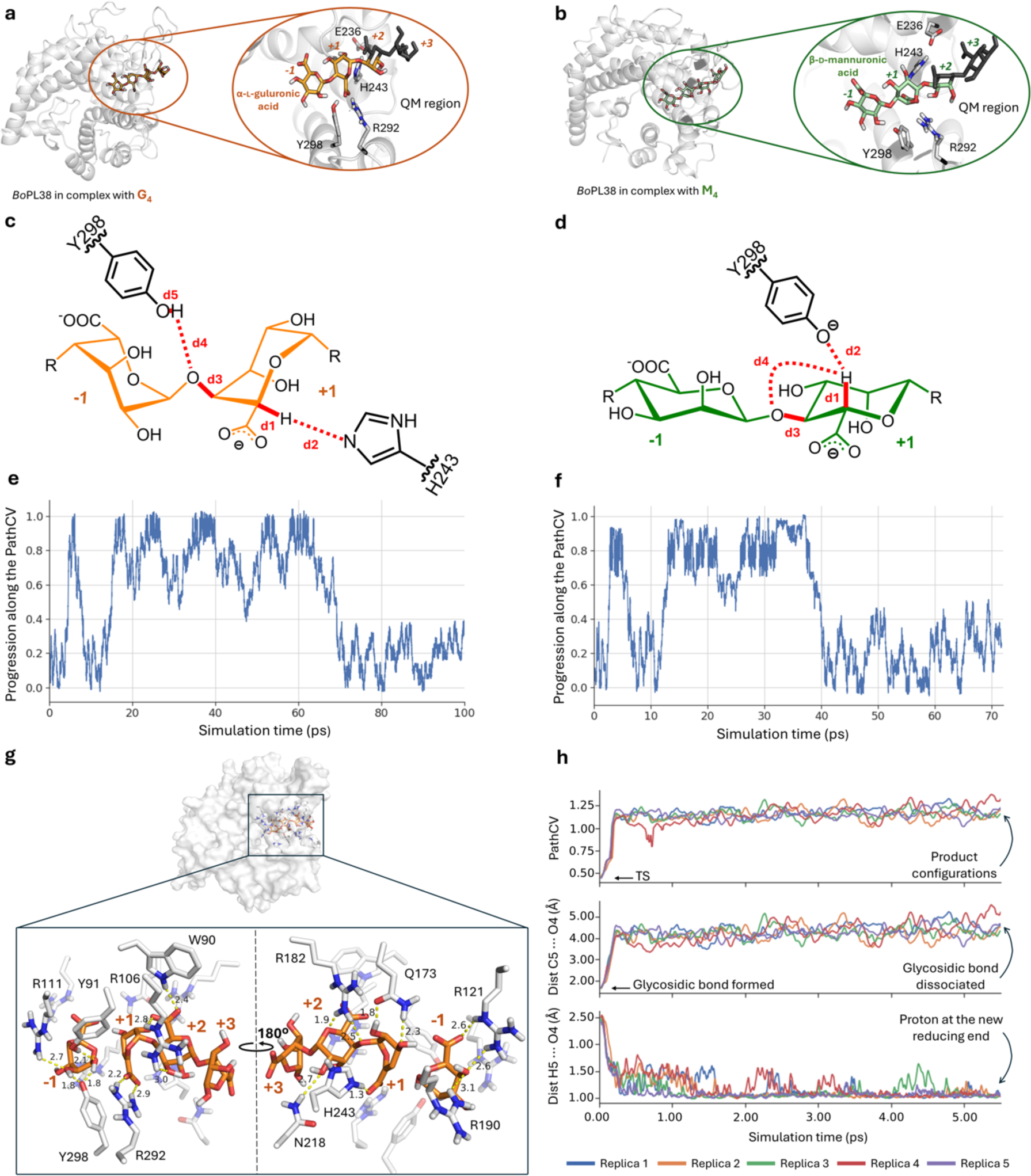
Structural insights from simulations of *Bo*PL38 reacting with G_4_ and M_4_. **a** Representative structural conformations during the QM/MM simulations, with a zoom-in highlighting the selected QM region for G_4_ and **b** M_4_. QM atoms shown in stick representation. Water molecules, and ions have been hidden for clarity. **c** Distances used to define the PathCV during the QM/MM OPES_E_ simulations of the *Bo*PL38 reaction on G_4_ and **d** M_4_, summarizing all proton movements along the reaction, and the cleavage of the glycosidic bond. **e** Progression along the PathCV during QM/MM OPES_E_ simulations of *Bo*PL38 in complex with G_4_ and **f** M_4_. Two recrossings were achieved during both simulations. **g** Extended view of the *Bo*PL38 active site post-catalysis of G_4_, showing G + Δ-GG following a QM/MM simulated reaction. Polar contacts are marked with yellow dashes, and the respective distances labeled in Å. **h** Unbiased QM/MM simulations of the TS described for the *syn* β-elimination reaction. Five replicas that ended up in product configurations were run for 5.5 ps. A quick change in the PathCV (top) is observed at the first stages of the runs, matching the fast cleavage of the glycosidic bond (middle). In all replicas, the proton was spontaneously transferred to the new reducing end after 5 ps (bottom).

**Extended Data Fig. 4.**
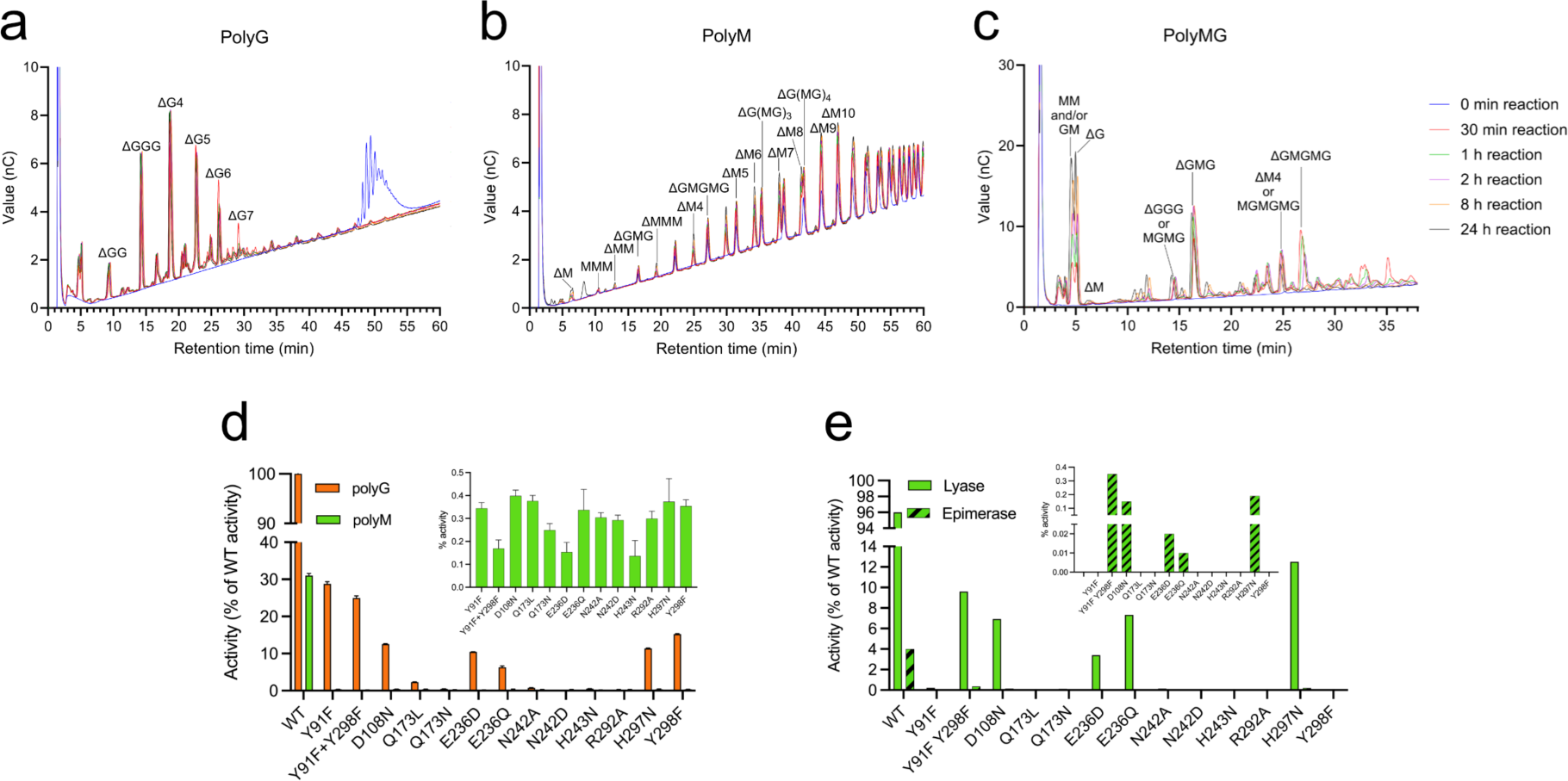
Product profile, reaction rates and lyase to epimerase ratio. **a-c** Overlaid time resolved HPAEC-PAD chromatograms of enzymatic degradation of 75 nM *Bo*PL38 on 0.5 mg ml^–1^ of **a** polyG, **b** polyM and **c** polyMG from 0 to 24 h. *Bo*PL38 degrades the substrates in a non-processive manner with products at the different time points getting shorter over time. Peaks are identified and marked from standards described in method section. **d** UV analysis of initial rates of *Bo*PL38 reactions on polyG (orange) and polyM (green). The insert highlights the much lower activity on polyM for all *Bo*Pl38 variants. **e** Lyase and epimerase activity of each *Bo*PL38 variant relative to the specific activity of the WT. The substrate used is ^13^C-1 labelled polyM. The lyase activity is based on the amount of substrate consumed during the reaction observed with time-resolved NMR, and the epimerase activity is based on the H/C-1 signals for M-G and Δ-G relative to other products formed in the HSQC recorded after the reaction was finished. The y-axis is discontinuous in panels **d** and **e**.

## Materials and methods

### Cloning, recombinant production, and purification

Genes encoding *B. ovatus* CP926 PL38 WT and mutants without signal peptide were purchased from GenScript (Piscataway, NJ, USA) and cloned into pET28a(+) using NheI and BamHI restriction sites in frame with an N-terminal His tag. Protein production and purification were performed using previously established methods^1^.

### Alginate substrates

PolyM (DP50–100 and DP6, F_G_ = 0.0), was obtained from an epimerase-negative AlgG mutant of *Pseudomonas fluorescens* as previously described^2^, polyG (DP10–30 and DP6, F_G_> 0.95) and polyMG (∼DP26 and DP6, F_G_ = 0.5) were prepared as previously described^3^ (Supplementary Fig. 6 and Supplementary Table 7). ^13^C-1 labeled polyM substrate was produced from the same *P. fluorescens* strain with ^13^C-1 labeled D-fructose as the carbon source. Δ-GG and Δ-MM trimers and Δ-GGG and Δ-MMM tetramers were prepared as previously described^4^.

### Specific activity and kinetics

The activity of ALs was determined spectrophotometrically (Bio-Tek Powerwave XS; Holm and Halby, Denmark) by following the formation of 4,5-unsaturated uronate units at 235 nm every 10 s at 37 °C. Enzyme and substrate were preincubated separately (5 min, 37 °C) before mixing in a 96-well UV-star chimney well plate (In Vitro, Australia). All measurements were performed in triplicate. Unless stated otherwise, the reaction conditions were 150 nM enzyme, 1 mg ml^–1^ substrate, 20 mM UB4 pH 7.5, and 200 mM NaCl. Protein concentrations were in all cases determined based on absorption at 280 nm and the molar extinction coefficient, calculated from the protein sequences, using the Expasy webserver (https://web.expasy.org/protparam/). Data are presented using GraphPad Prism 10, version 10.2.2.

### NMR spectroscopy (general sugar identification and epimerase activity)

For the time-resolved NMR analysis of reactions, a stock solution of substrate in 10 mM HEPES, pH 7.5 with 200 mM NaCl in 99.9% D_2_O (Sigma-Aldrich) was prepared. Substrate stock (150–200 μl) solution was transferred to a 3 mm LabScape Stream NMR tube (Bruker LabScape). A 1D ^13^C spectrum was recorded prior to the time-resolved NMR experiment to ensure that the sample had not undergone any degradation or contamination. The enzyme was added to the preheated substrate and mixed by inverting the sample several times. The sample was inserted into the preheated NMR instrument and the measurement was started. A 1D ^13^C (zgig30) NMR spectrum was recorded every 2, 5, or 10 min, resulting in a pseudo-2D spectrum. Sample and acquisition parameters are given in Supplementary Tables 5–6. After completion of the time-resolved experiment, a ^1^H-^13^C HSQC (heteronuclear single quantum coherence with multiplicity editing, hsqcedetgpsisp2.3) spectrum was recorded.

In-depth characterization of the reaction products of *Bo*PL38 with three different substrates was performed by mixing the enzyme (1.85 µM) with 200 µl of 9.4 mg ml^–1^ substrate in 10 mM HEPES, pH 7.5 with 200 mM NaCl in 99.9% D_2_O and letting the reaction proceed at room temperature for 24 h. The substrates were polyM (DP50–100), polyMG (∼DP26), and polyG (DP10–30). After 24 h reaction, the mixture was moved to a 3 mm NMR tube, and the following spectra recorded for assignment of reaction products formed: 1D proton with water suppression (noesygppr1d), ^1^H-^13^C HSQC, ^1^H-^13^C HMBC (heteronuclear multiple bond coherence with suppression of one-bond correlations, hmbcetgpl3nd), ^1^H-^1^H IP-COSY (in-phase correlation spectroscopy, ipcosyesgp-tr)^5^.

Δ-GG (4.8 mg ml^–1^) and Δ-MM (5.0 mg ml^–1^) trimers were dissolved in 99.9 % D_2_O with TSP 0.0125 w/v% for internal referencing. ^1^H 1D, ^1^H-^13^C HSQC, ^1^H-^13^C H2BC (heteronuclear two bond correlation spectroscopy, h2bcetgpl3pr), HMBC, IP-COSY, and ^1^H-^1^H TOCSY (total correlation spectroscopy, mlevphpr) spectra were recorded to allow full assignment of the resonances of the trimers.

The purity of the substrates was confirmed using NMR spectroscopy by recording a ^1^H 1D and ^1^H-^13^C HSQC spectrum of each of polyMG, polyG, and polyM. Resonances were assigned based on the assignment of the two trimers and references^6^. The ^1^H-^13^C HSQC spectra are shown in Supplementary Fig. 6 and the assignment in Supplementary Table 7.

All NMR spectra were acquired at 25 °C on a Bruker Ascend 800 MHz spectrometer or a Bruker Ultrashield II 600 MHz spectrometer both with Avance III HD console using a 5 mm Z-gradient CP-TCl (H/C/N) cryogenic probe. Both the 800 MHz and 600 MHz NMR spectrometers were located at the NV-NMR-Center/Norwegian University of Science and Technology (NTNU). ^1^H signals were internally referenced to the water signal and ^13^C signals were indirectly referenced to the water signal based on absolute frequency ratios^7^ except for when otherwise described. The spectra were recorded using TopSpin version 3.5 and processed and analyzed using TopSpin version 4.3.0 (Bruker BioSpin AG).

Key signals were integrated in the time-resolved reaction spectra to monitor the reaction progress. All integrals in all spectra were normalized to the total ^1^H integral from the last obtained spectra (%).

Time-resolved NMR was performed with ^13^C-1 labeled polyM and *Bo*PL38 WT or mutants, and afterward, HSQC was acquired (Supplementary Tables 8–9). The activity of each *Bo*PL38 WT and mutant was calculated based on time-resolved NMR signals by integrating all C-1 signals, normalizing the total integrals, extracting the integral of the <u>M</u>-M substrate C-1 signal (refers to the underscored) relative to the total integral in the final spectra and calculated as a percentage of WT activity. Epimerase activity was measured as the integral of the H\C-1 signals for <u>M</u>-G and Δ-<u>G</u> in the HSQC relative to the total integral of H\C-1 signals (refers to the underscored), converted to a percentage of the total activity of the mutant, and adjusted for concentration difference from the WT. Lyase activity was defined as the rest of the total activity when epimerase activity was subtracted, and it was also adjusted for concentration difference from the WT.

### High-performance anion-exchange chromatography (HPAEC) with pulsed electrochemical detector (PAD)

Model substrates (0.5 mg ml^–1^) of alginate, polyG (DP25–30), polyM (∼DP27) and polyMG (DP24), were incubated with 75 nM *Bo*PL38 in 20 mM HEPES, 150 mM NaCl, pH 7.5 at 37 °C. Samples were taken before the addition of enzyme (t= 0 min) and at 30 min, 1 h, 2 h, 8 h, and 24 h. Each sample was inactivated by incubating for 10 min at 90 °C, centrifuged at 10,000 rpm, and the supernatant stored at −20 °C until analysis on an ICS-5000+ system (Thermo Scientific) with IonPac 4 x 250 mm IonPac AS4A and 4 x 50 mm AG4A guard columns (Thermo Scientific). Samples were eluted at 24 °C with 0.1 M NaOH (isocratic) and a sodium acetate gradient of 8.75 mM min^–1^, starting at 10 mM sodium acetate, with a flow rate of 1 ml min^–1^ over 90 min. Waveform A (Gold-Ag-AgCl Re, Carbo, quad) was used for detection. Data were collected and processed with Chromeleon (Thermo Scientific) 7.2 software and presented using GraphPad Prism 10, version 10.2.2. Standards used for HPAEC-PAD analysis were polyMG (F_G_ = 0.46), partially degraded by the M-lyases from *Haliotis tuberculata* (*H.tub*)^8^ and from *Azotobacter vinelandii* (AlgL)^9^ breaking G-M and M-G bonds, respectively. PolyM (F_G_ = 0.00) was partially degraded by the *H.tub.* M-lyase and polyG (F_G_ >0.97) was partially degraded by a G-lyase (AlyA) from *K. pneumoniae*^3^. Moreover, the three substrates were subjected to acid hydrolysis under conditions only yielding saturated oligosaccharides^10^.

### Nano differential scanning fluorimetry

Thermal shift assays were performed on a Prometheus Panta (Nanotemper) instrument. Protein concentrations were adjusted to 2 µM in 20 mM HEPES pH 7.5, and 150 mM NaCl before being loaded into glass capillaries. Measurements were carried out in triplicate. Subsequent analysis of T_M_ was performed the Panta Analysis software suite (Nanotemper).

### Crystallization, diffraction data collection and processing

Initial crystals of *Bo*PL38 WT (8.4 mg ml^–1^) in 20 mM HEPES pH 7.5 and 150 mM NaCl were grown by sitting-drop vapor diffusion in SWISS-CI MRC-3 drop plates (Molecular Dimensions) by mixing 0.2 µl enzyme with 0.1 µl reservoir solution consisting of 25% (w/v) PEG3350, 0.2 M (NH_4_)_2_SO_4_ and 0.1 M BIS-TRIS pH 5.5. Optimized crystals of both *Bo*PL38 WT and mutants were grown by sitting-drop vapor diffusion in MRC-MAXI plates (Molecular Dimensions) with 2 µl enzyme solution (8.4 mg ml^–1^) against 1 µl reservoir consisting of 14–18% (w/v) PEG3350, 0.2 M (NH_4_)_2_SO_4_ and 0.1 M citric acid pH 3.5 or 0.1 M HEPES pH 8.0.

Crystallographic complexes were obtained by dissolving a few AOS powder grains in 3 µl of the reservoir solution and adding to the drop containing an already-grown crystal. The AOS powder was either purified hexa-β-D-mannuronic acid (M_6_), tetra-β-D-mannuronate (M_4_), hexa-⍺-L-guluronate (G_6_), tetra-*α*-L-guluronate (G_4_), mixed MG residues at DP6 ((MG)_3_), or unsaturated tetramers of G or M (Δ-GGG, Δ-MMM). The protein crystals were allowed to equilibrate with the dissolved AOS for 2 h at RT, then cryoprotected with 20% glycerol in reservoir solution for 10 s and cryocooled in liquid nitrogen. Data were collected at BioMAX at the MAX IV Laboratory^11^, using MXCube3^12^ data collection software and IspyB/Exi monitoring and management software^13^. The data was processed and scaled using XDS/XSCALE^14^ (version Jan 10, 2022) or through xia2 (3.5.0) (Supplementary Table 1 for data collection statistics).

### Structure phasing, refinement, and analysis

Structures of *Bo*PL38 WT and mutants binding AOS were determined using the phases of the original *Bo*PL38 WT structure (PDB 5BDQ)^1^ as the input model in initial rigid body refinement using phenix.refine (1.21_5207), or through molecular replacement using PHASER (1.21_5207). Subsequent refinement was done by alternating restrained refinement in phenix.refine and manual rebuilding and validation in COOT ^15^ (0.9.8.92). The AOSs were built into the available electron density within the active site of all 4 molecules of the asymmetric unit, and the AOS geometry was validated using Privateer^16^ (version MKIV: 06/02/21) in the CCP4 suite^17^ (8.0.017) with the CCP4i2 (1.1.0 revision 6539) interface, and additionally analyzed in PyMOL (Schrödinger, 2.1). Further validation of protein and sugar geometry was performed using MolProbity^18^ (4.5.2) in the phenix suite.

### Classical MD simulations of enzyme complexes

Crystal structures of *Bo*PL38 in complex with G_4_ and M_4_ at pH 8 (PDB 9FHW and 9FHX) were used to conduct all molecular dynamics (MD) reported in this article. Simulations were performed under conditions of neutral pH, i.e. with all arginine and lysine residues taken as positively charged, while all aspartate and glutamate residues, as well as all carboxylate groups of the M and G moieties, were taken as negatively charged. The protonation state of histidine residues was chosen according to their computed p*K*_a_ values using the H++ server^19–21^. Since H243 and Y298 can easily exchange a proton via water molecules (Supplementary Fig. 17), their protonation state was interchanged according to their catalytic roles in G and M complexes. Each system was enclosed in a periodically repeated rectangular box of sizes 77.725 x 83.850 x 90.199 Å^3^ and 77.725 x 83.850 x 90.199 Å^3^ in G and M complexes, respectively, allowing a minimum distance of 15 Å between the solute surface and the box edges. Boxes were solvated with TIP3P^22^ water molecules, and sodium and chlorine ions were added to ensure system neutrality. The protein was described using the Amber ff14SB force field^23^, while the GLYCAM06 force field^24^ was used to describe the carbohydrate moieties. In the M complex simulations, the atomic partial charges (RESP) of Y298 (negatively charged) were calculated at the HF/6-31G* level theory using Gaussian09^25^. The topology and coordinate files for the classical MD (cMD) simulations were generated using the LEaP module of AmberTools21^26^. All cMD simulations were run using Amber20^26^ with the CUDA version of the PMEMD module^27^. The energy of the solvated system was minimized with a three-step protocol (minimization of solvent, solvent plus ligand, and finally the whole system). Each step in the protocol involved 5000 steps of steepest descent and 5000 steps of conjugate gradient minimization. Subsequently, the system was gradually heated to 308 K in the NVT ensemble using the Berendsen thermostat, with positional restraints applied to the heavy atoms of both the protein and the ligand. The restraints were gradually decreased over three 500 ps steps, followed by 1000 ps of NPT equilibration using the Berendsen thermostat and barostat to maintain the system at 308 K and 1 atm for density equilibration. Positional restraints were applied to distances between the ligand and the protein as well as in dihedral angles controlling the conformation of the sugar at subsite +1 in a three-step protocol decreasing gradually the force constants (100 to 20 to 5 kcal mol⁻¹) to ensure the relaxation of the system.

Restraints were released after 20 ns of simulations, keeping just one restraint on the sugar conformation to avoid the preference for chair conformations in empirical force fields, as previously described^28^. Three replicas of 500 ns were considered, taking 100 ns as equilibration and 400 ns as production (Extended data Fig. 2). The SHAKE algorithm^29^ was used to constraint all bonds involving hydrogen atoms and the MD time step was set to 2 fs. Production runs in the NPT ensemble were controlled by the Langevin thermostat and the Berendsen barostat.

### QM/MM modelling of the β-elimination reaction

QM/MM MD simulations were used to study the reaction mechanism. The QM region was treated by DFT-based MD, while the MM region was described using cMD. A representative snapshot from the cMD trajectories of *Bo*PL38 in complex with M_4_ and G_4_ were used as a starting point for running two independent QM/MM MD simulations. In both cases, the QM region accounted for 103 atoms, including residues E236, H243, R292, and Y298 (with QM/MM border located at the Cβ-Cα bond), and sugars at subsites –1, +1 and some atoms at subsite +2 (Extended data Fig. 3a–b). The QM box size was 20.17 x 18.52 x 26.82 Å^3^ in the G complex while 24.09 x 20.87 x 19.69 Å^3^ in the M complex. The QM atoms at the QM/MM border were saturated with hydrogen atoms using the IMMOM approach^30^. The CP2K v9.0^31^ program was used for the QM/MM MD simulations. The QM region was described at the DFT (PBE) level of theory, using the dual basis set of Gaussians and plane-waves (GPW) formalism. The wave function was expanded by a Gaussian triple-ζ valence polarized (TZV2P) basis set, and the electron density was converged using the auxiliary plane wave basis set with a density cut-off of 330 Ry and GTH pseudopotentials^32^. After geometry optimization to ensure the relaxation of the QM region, the system was equilibrated by QM/MM MD for 10 ps, before starting enhanced sampling simulations of the chemical reaction. The on-the-fly probability enhanced sampling (OPES) method^33,34^ in its exploratory variant (OPES_E_) was used to explore the Free Energy Surface (FES). OPES is a newly developed method that shares features with metadynamics^35^ and has been proven to be very efficient in obtaining reaction free energy landscapes complex processes, including enzyme reactions^36^. To activate the chemical reaction and drive the systems from reactants to products, a PathCV^37,38^ was designed based on the main interatomic distances involved in the hypothesized mechanism (Extended data Fig. 3c–d). The initial guess of the path was set from prior modeling of the β-elimination mechanisms of a PL7 lyase^36^, as well as preliminary OPES_E_ simulations of the systems using ‘canonical/classical’ CVs (data not shown). Only the *s* component of the path (progression along the path) was biased, while a potential wall was put on the z component (distance of the system with respect to the defined path) at 0.8 with a force constant of 500 kcal mol⁻¹, allowing the system to visit other possible pathways without being restricted to the one described. The OPES_E_ engine was set up with a PACE of 100, an Adaptive Sigma stride of 200, and a barrier parameter of 20 kcal mol⁻¹. The use of those parameters allowed us to have a better description of the TS, avoiding the visiting of high-energy regions. Simulations were stopped after two recrossing over the TS (Extended data Fig. 3e–f), and the FES was computed using a reweighting scheme. The transition states were further refined using an iso-committor analysis (Supplementary Figs. 18–19). The computed free energy barriers were compared with experimental values estimated using Transition State Theory and the Eyring equation^39^, based on the measured rate constants for the enzymatic reactions. The experimental rate constants were 9.1 ± 0.11 s⁻¹ for polyG and 0.9 ± 0.02 s⁻¹ for polyM^1^, resulting in the experimental free energy barriers 16.8 kcal mol⁻¹ for polyG and 18.2 kcal mol⁻¹ for polyM. All QM/MM MD simulations were performed in the NVT *ensemble* using a coupling constant of 10 fs and an integration time step of 0.5 fs. The trajectories from the simulations were visualized using VMD^40^ and analyzed using VMD, the driver module of PLUMED^41^, and the cpptraj module^26^ of AMBER. All plots were generated using the MatPlotlib^42^ package of python and figures of the structures were generated and rendered using PyMOL (Schrödinger, 2.1).

*Classical MD simulations of BoPL38 WT and variants.* All the mutants were designed *in silico* by replacing the residues using PyMOL (Schrödinger, 2.1). The systems were set up following the same scheme as the one described above (*Classical MD simulations of the complexes*) and excluding the parts related to the ligand (simulations were run on the plain enzymes). Three replicas were run for each system WT, D108N, and R292A at 308 K in the NPT ensemble, for 200 ns each and taking the first 50 ns as equilibration (Supplementary Fig. 13), while one replica was run for the H243D and N242D variants (Supplementary Fig. 11a and d).

