## Supplementary figures and tables for "The Swiss Army Knife of Alginate Manipulation – A Gut Bacterium Alginate Lyase with Diverse Catalytic Activities"

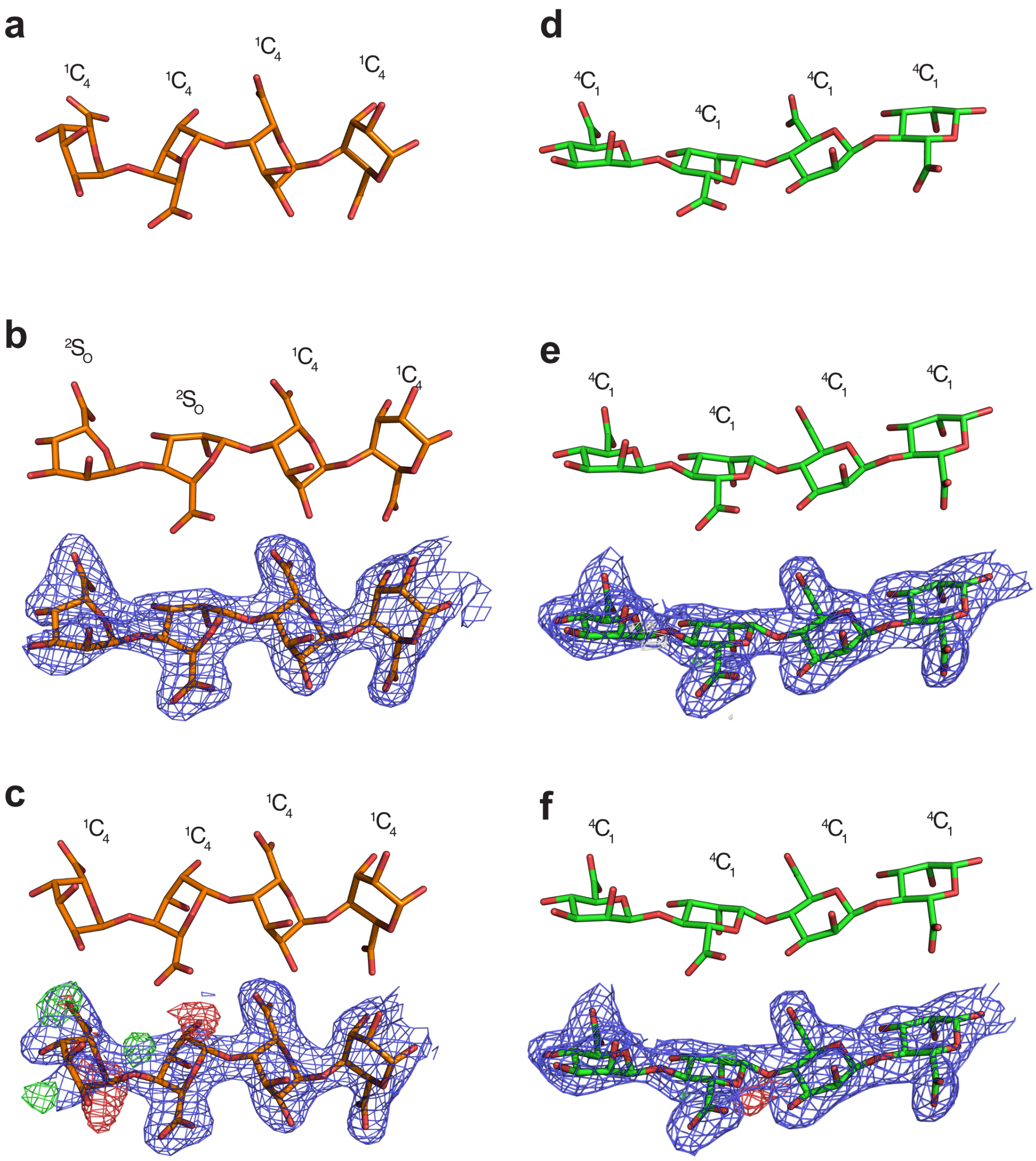

**Supplementary Fig 1. | Fitting substrates into the active site of *Bo*PL38**. The uronic acid units are shown from three stages of active site modelling. **a** Computationally relaxed and unrefined $G_4$. **b** Geometrically restrained $G_4$ after refinement. No difference density visible. **c** $G_4$ after refinement with increased geometry restraints. The geometry at subsites –1 and +1 is restrained to the $^1C_4$ conformation. However, the difference density suggests that the O2 and O3 atoms of C2 and C3 OH groups at subsite –1 should be reorientated to adopt the $^2S_0$ conformation. **d** Computationally relaxed $M_4$. **e** $M_4$ after geometrically restrained refinement. No difference density visible. **f** $M_4$ after refinement with increased geometrical restraints. M in subsite +1 is in the $^4C_1$ conformation. The negative difference density suggests this geometry should be slightly distorted towards $^2S_O$. In panels B, C, E, and F the sugars are shown with and without $2F_O$-$F_C$ electron density at 1.0 σ contour level (in blue) from the refinement and, where present, the $F_O$-$F_C$ difference density in green and red, for ±3.0 σ.

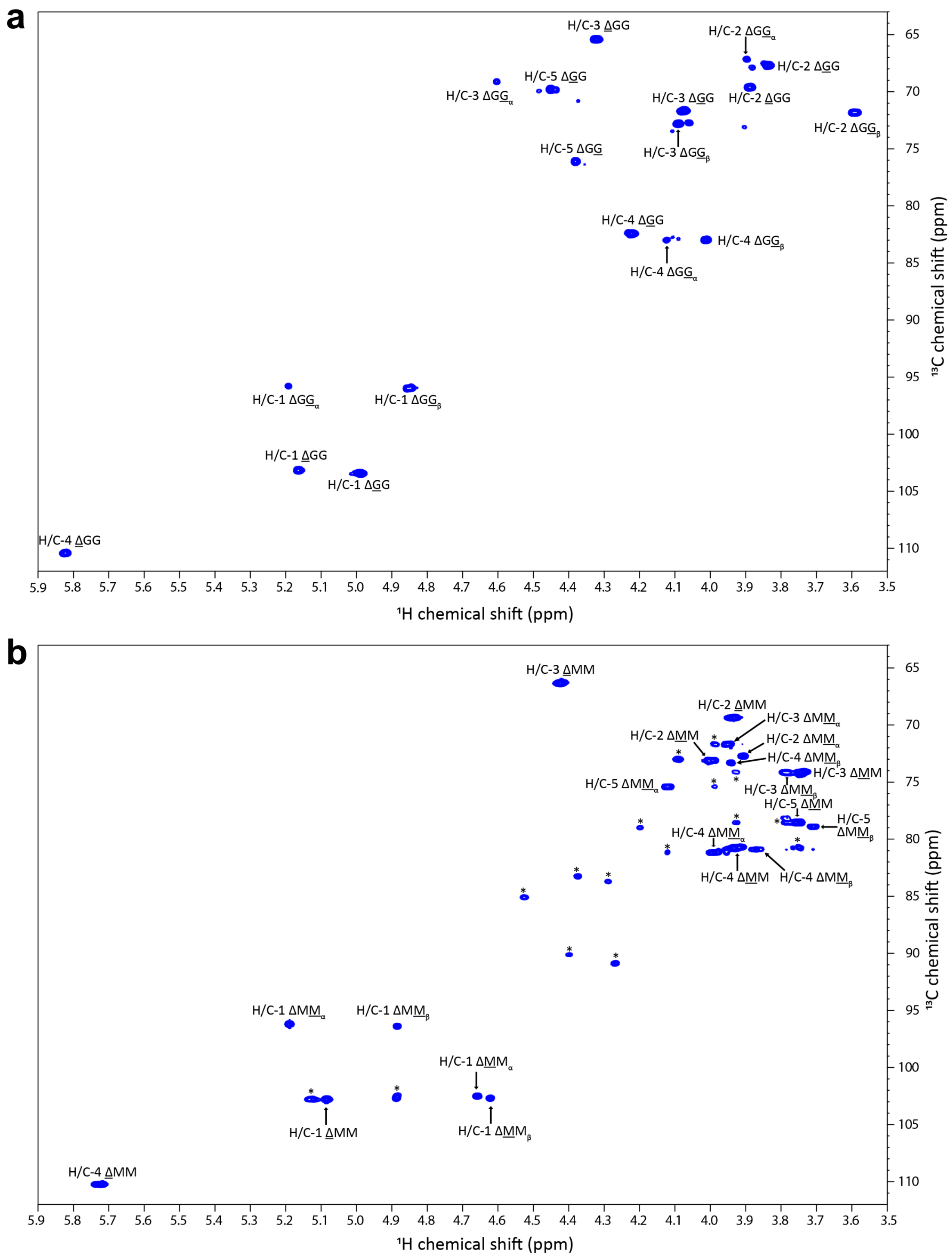

**Supplementary Fig. 2 |** $^{1}H$-$^{13}C$ HSQC of **a** Δ-GG trimer (4.8 mg ml$^{-1}$) and **b** Δ-MM trimer (5.0 mg ml$^{-1}$), both at pH 7 and 25 °C with TSP 0.0125 w% in 99.9 % $D_2O$. All resonances were assigned using $^{1}H$-$^{13}C$ HSQC, H2BC, HMBC, IP-COSY, and TOCSY spectra. An impurity (signals marked with *) is present due to Schiff-base formation during processing of the Δ-MM trimer.

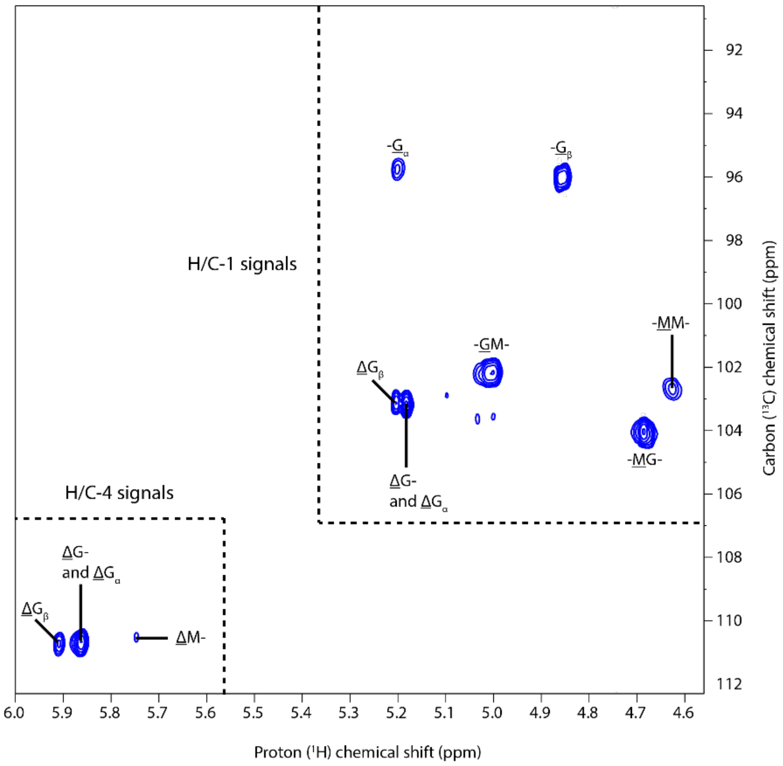

**Supplementary Fig. 3 |** Anomeric and Δ H/C-4 regions of $^{1}H$-$^{13}C$ HSQC of polyMG (10.0 mg $ml^{-1}$) treated with *Bo*PL38 WT (413 nM) for 13 h at pH 7.5 and 25 °C with HEPES 10 mM, NaCl 200 mM in 99.9% $D_2O$.

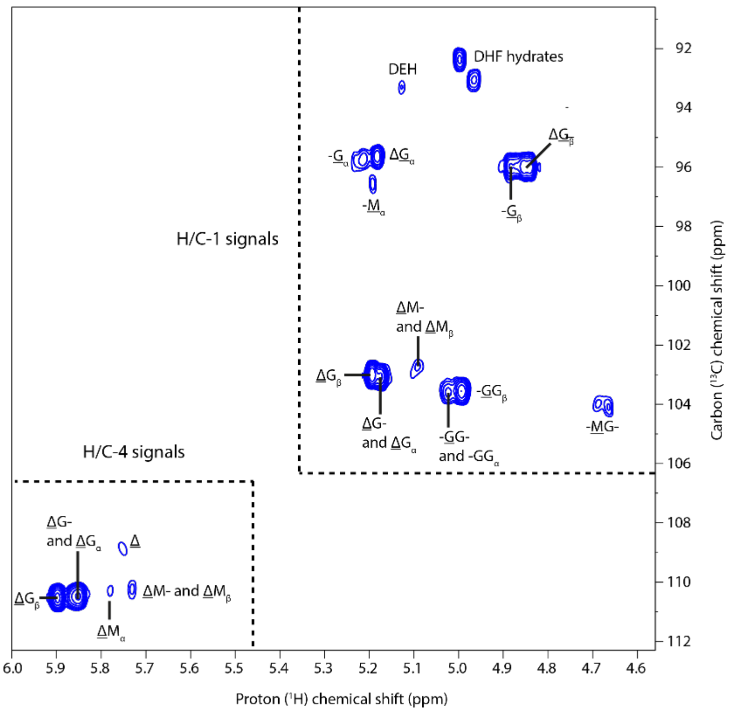

**Supplementary Fig. 4 |** Anomeric and Δ H/C-4 regions of $^{1}$H-$^{13}$C HSQC of polyG (9.0 mg ml$^{-1}$) treated with *Bo*PL38 WT (413 nM) for 13 h at pH 7.5 and 25 °C with HEPES 10 mM, NaCl 200 mM in 99.9% $D_2O$.

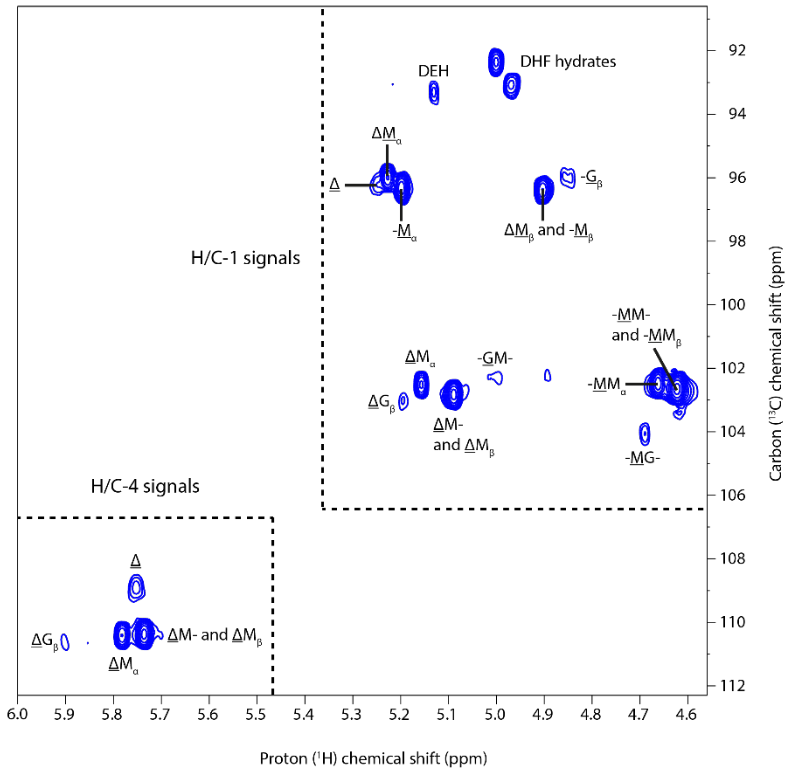

**Supplementary Fig. 5 |** Anomeric and Δ H/C-4 regions of $^{1}H$-$^{13}C$ HSQC of polyM (10.0 mg ml$^{-1}$) treated with *Bo*PL38 WT (413 nM) for 13 h at pH 7.5 and 25 °C with HEPES 10 mM, NaCl 200 mM in 99.9% $D_2O$.

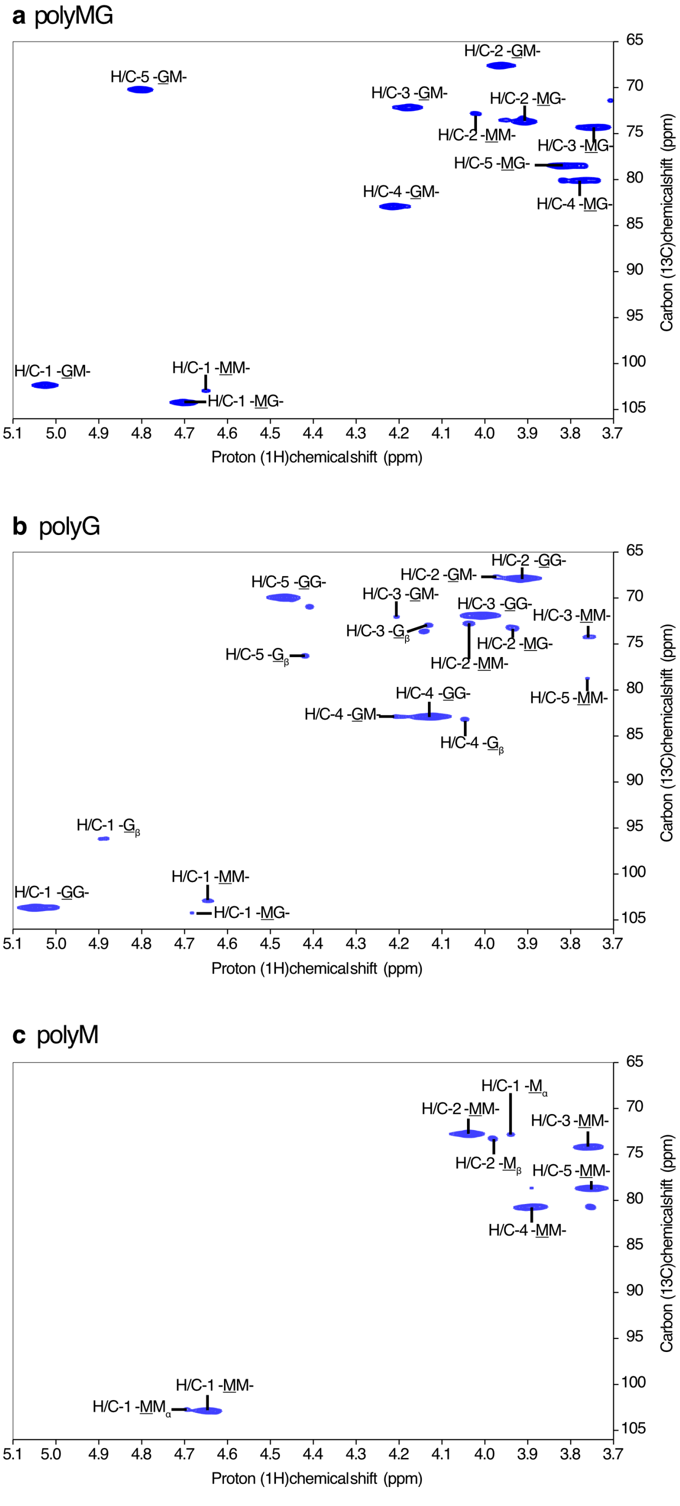

**Supplementary Fig. 6 | Substrate purity.** $^{1}$H-$^{13}$C HSQC of three substrates used for NMR. All NMR spectra were recorded at 25 °C with TSP 0.0014 w% in 99.9 % $D_2O$. **a** polyMG (DP 24, $F_G$ = 0.5, 10 mg mL$^{-1}$), **b** polyG (DP 10-30, $F_G$ > 95, 10 mg/mL$^{-1}$), **c** polyM (DP 50-100, $F_G$ = 0.0, 7.0 mg/mL$^{-1}$). The spectra show that polyMG and polyG contain minor MM signals, while polyM contains no MG or GG signals. Assignment performed based on assignments shown in S12–S13 and Holtan *et al.* 2006[1].

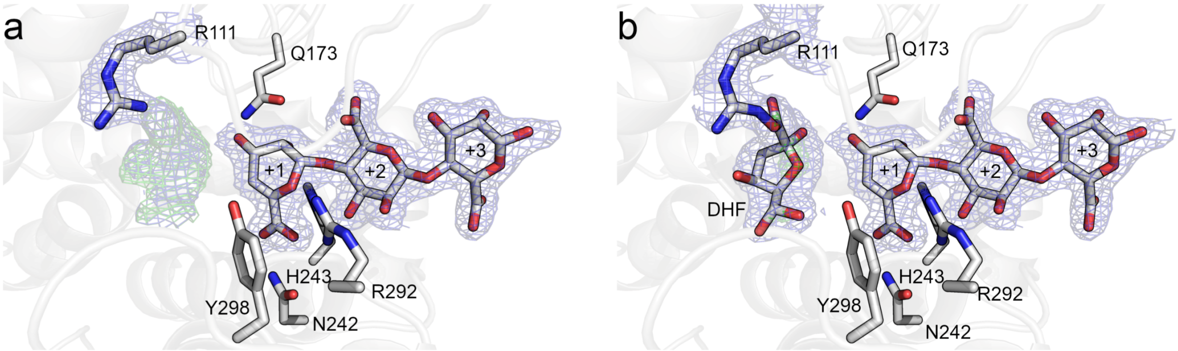

**Supplementary Fig. 7 | *Bo*PL38 complex with unsaturated non-reducing end Δ-M$_3$.** Soaking the tetrasaccharide Δ-M$_3$ into crystals of *Bo*PL38-WT resulted in a clearly defined and bound Δ-M$_2$ substrate. Similar results are obtained with Δ-G$_3$ soaked crystals (not shown). **a** The active site contained a cloud of electron density and difference density, which indicated a missing component in the model. **b** Placing in the downstream product DHF appears to satisfy the density. However, the modeled conformation clashes with R111. For this reason, the final structures of both *Bo*PL38-ΔM$_3$ and *Bo*PL38-ΔG$_3$ do not contain DHF and the cloud of electron density has been left unmodeled. Electron density is displayed as 2F$_O$-F$_C$ in blue mesh at 1.0 σ. Difference density is shown as F$_O$-F$_C$ in green and red mesh at ±3.0 σ.

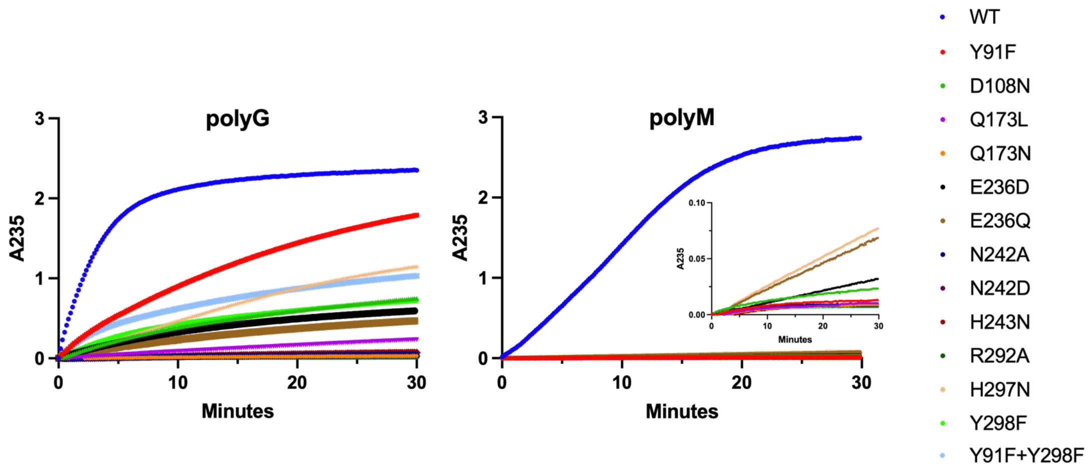

**Supplementary Fig. 8 | UV analysis of *Bo*PL38 WT and variants reacting on polyG and polyM.** Progression curves for *Bo*PL38 reacting on polyG and polyM. Insert in polyM panel highlights the much lower activity for *Bo*PL38 variants reacting with polyM. In most cases, any mutation applied to the *Bo*PL38 enzyme will abolish the activity on polyM, while for polyG, some activity is retained.

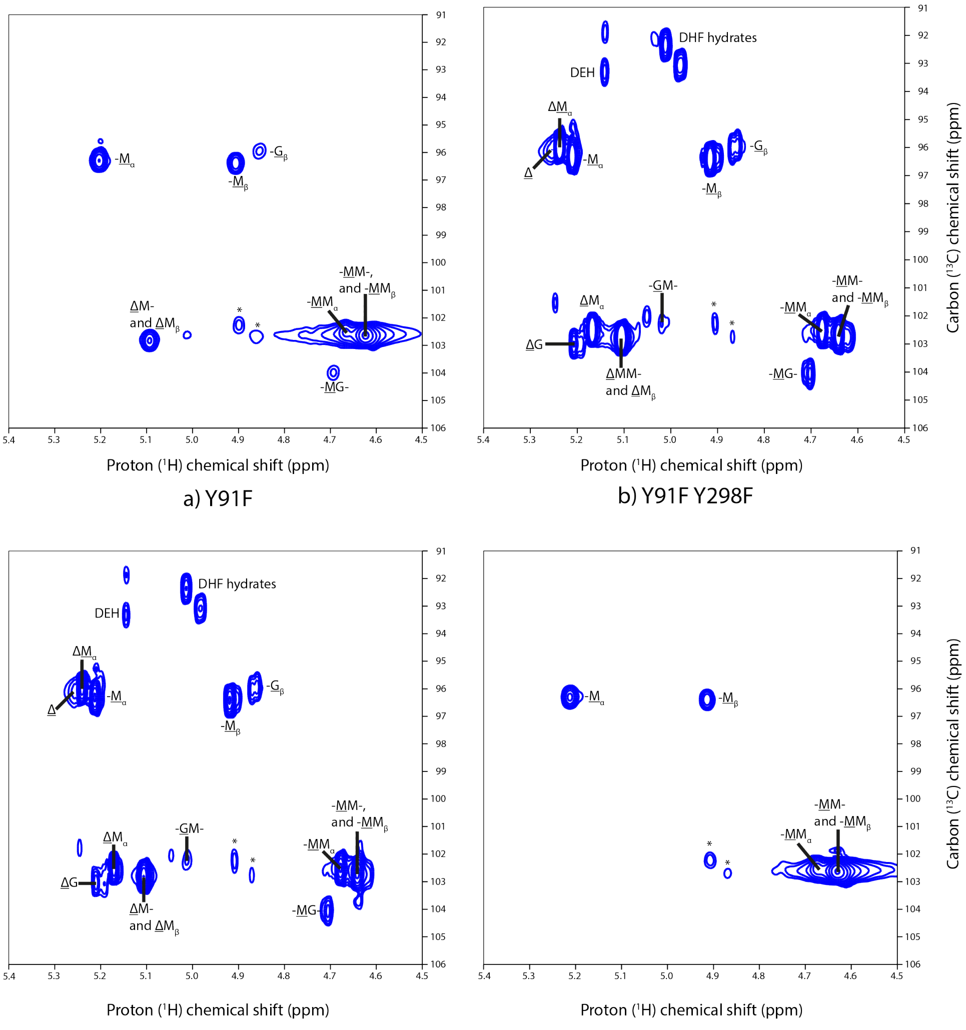

a) Y91F

b) Y91F Y298F

c) D108N

d) Q173L

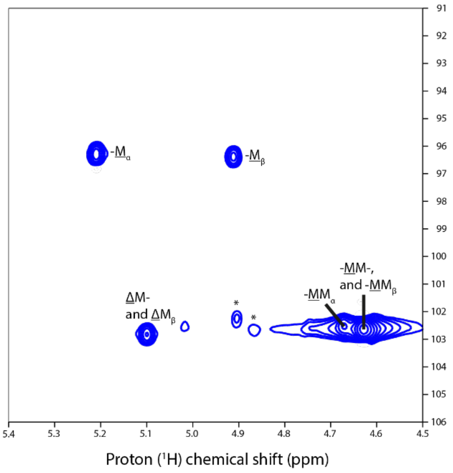

e) Q173N

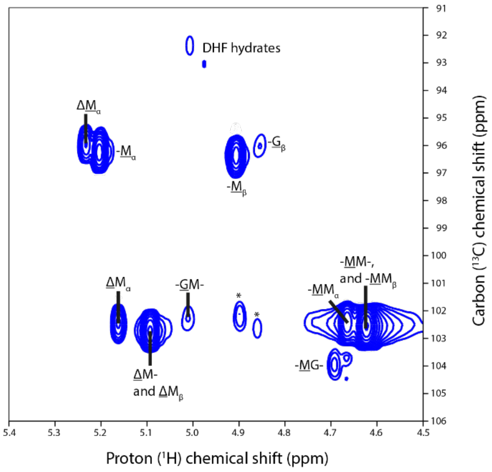

f) E236D

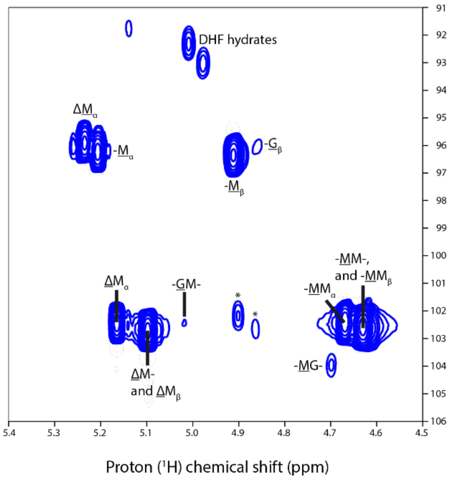

g) E236Q

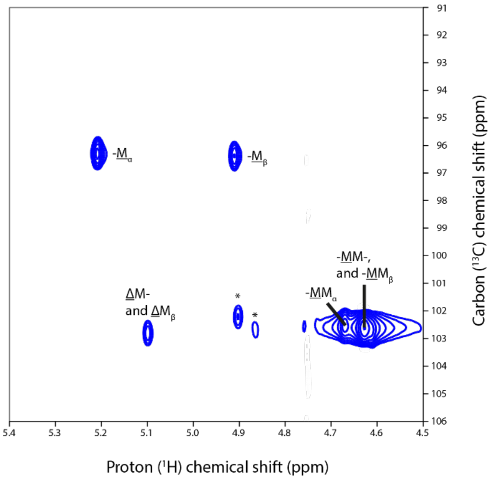

h) N242A

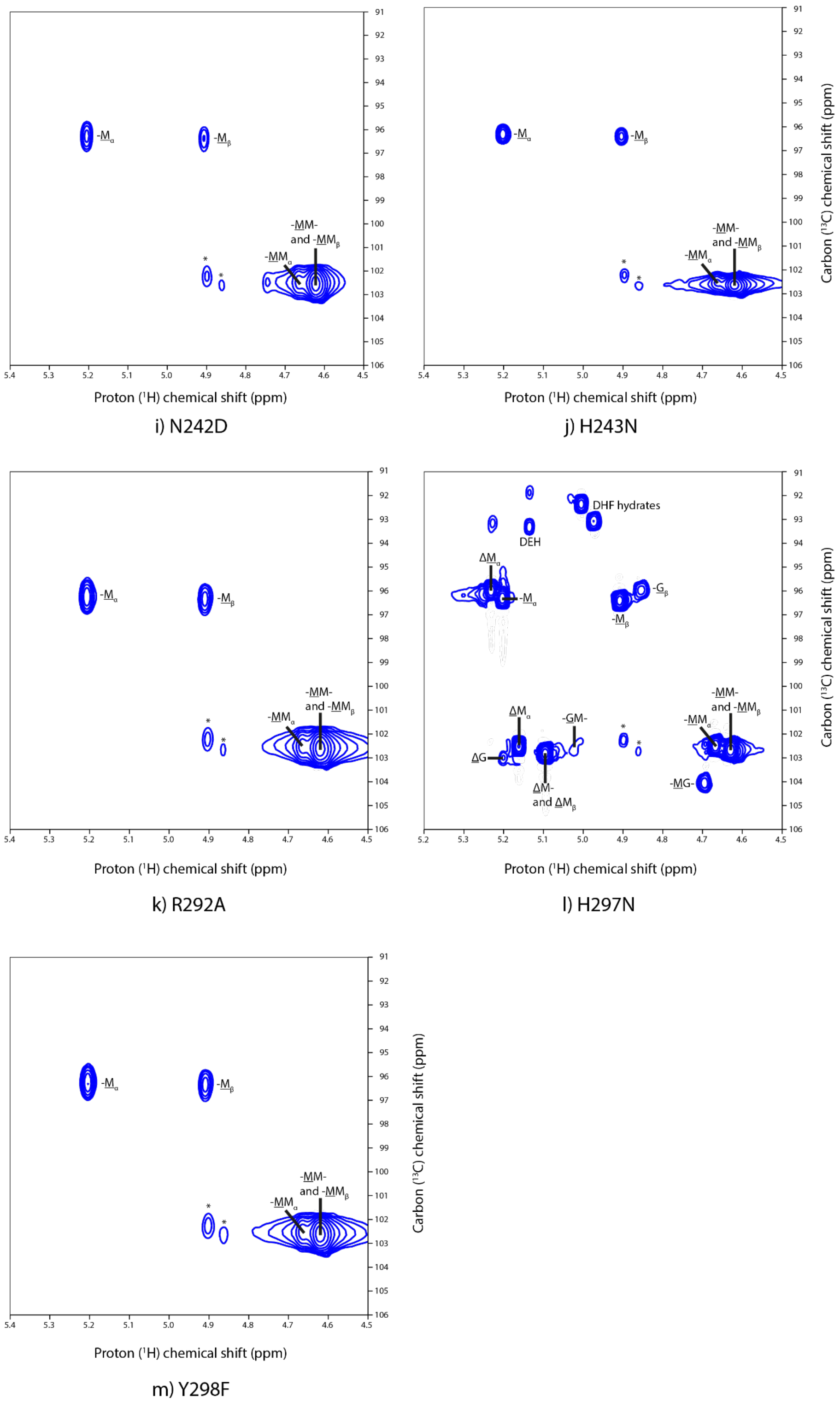

**Supplementary Fig. 9 |** Anomeric region of $^{1}$H-$^{13}$C HSQC of polyM (7.0–9.0 mg ml$^{-1}$) treated with individual *Bo*PL38 variants (3.65–6.95 μM) for 10–12 h at pH 7.5 and 25 °C with HEPES 10 mM, NaCl 200 mM in 99.9% $D_2O$. Except *Bo*PL38 N242A (h) for which the reaction only proceeded for 6.5 h. An impurity (marked with *) is present in all reaction mixtures, and the HSQC spectra are scaled so that the impurity signals have the same intensity in all spectra.

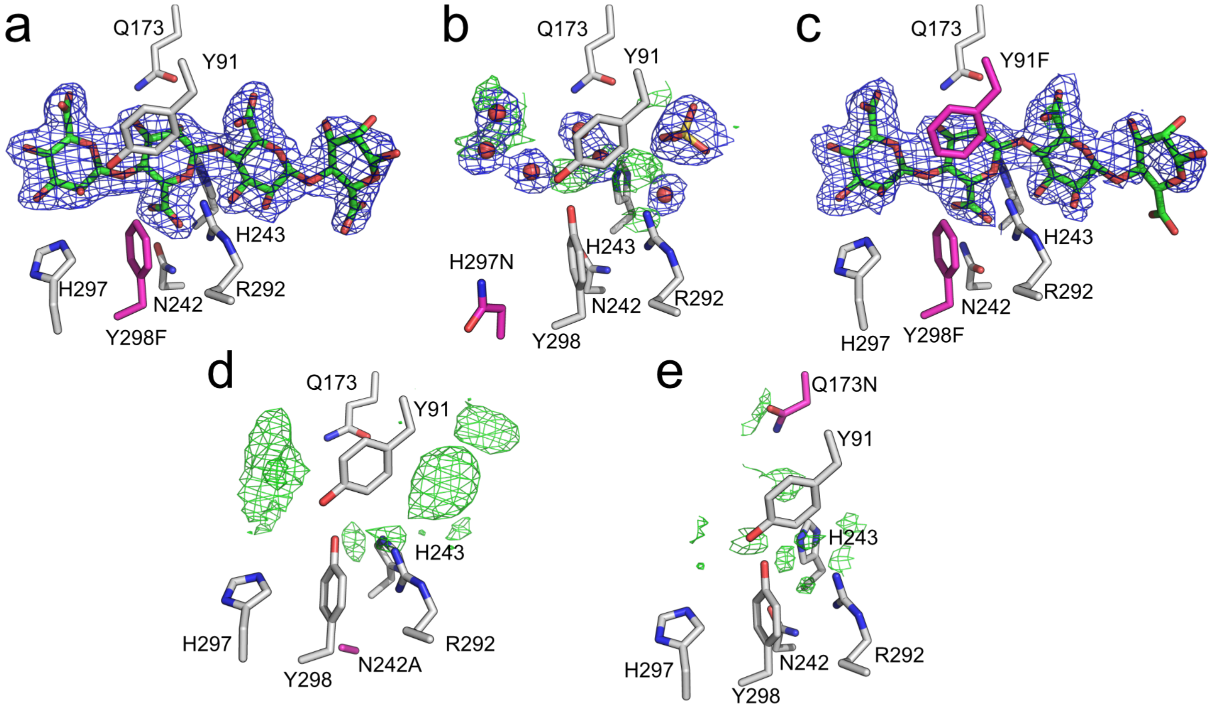

**Supplementary Fig. 10 | Active site structures of *Bo*PL38 mutants.** Crystal structures of variants of *Bo*PL38 elucidate the functional role of different amino acid residues. **a** Y298F retains the ability to bind $M_4$ *in crystallo*, and Y298 is therefore not required to form the Michaelis-complex, although activity is lost. **b** H297N is unable to form complex with $M_4$ *in crystallo* and H297 is therefore required to do so. **c** The double mutant Y91F+Y298F retains $M_4$ binding *in crystallo* and some activity. **d** N242A and **e** Q173N lose activity and $M_4$ binding *in crystallo* and are main contributors in binding sugars at subsite +1. Electron density, $2F_O$-$F_C$, shown in blue mesh at 1.0 σ. Difference density, $F_O$-$F_C$, shown in green and red mesh at ± 3.0 σ.

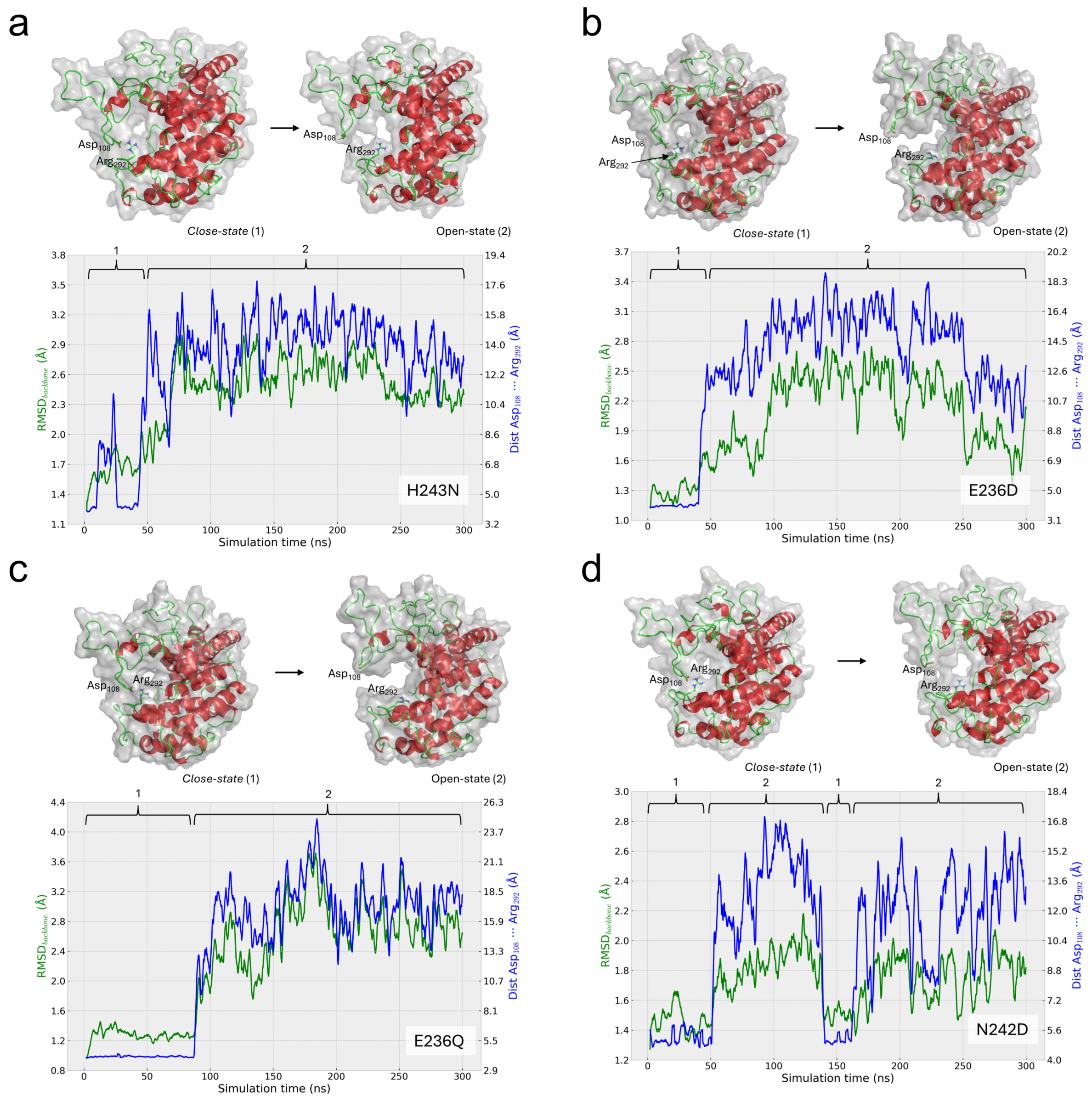

**Supplementary Fig. 11 | Stability analysis of *Bo*PL38 variants H243N, E236D, E236Q and N242D classical MD simulations.** Structure representations over the course of the 300 ns simulation are shown. **a** The model containing the H243N mutation becomes destabilized after 40 ns. **b** The model containing the E236D mutation becomes destabilized after 45 ns. **c** The E236Q variant has a significant shift in RMSD at 90 ns of the simulation. **d** The N242D variant has a significant shift in RMSD at 50 ns of the simulation. These observations together with the distance between R292 and D108 indicate the point of destabilization.

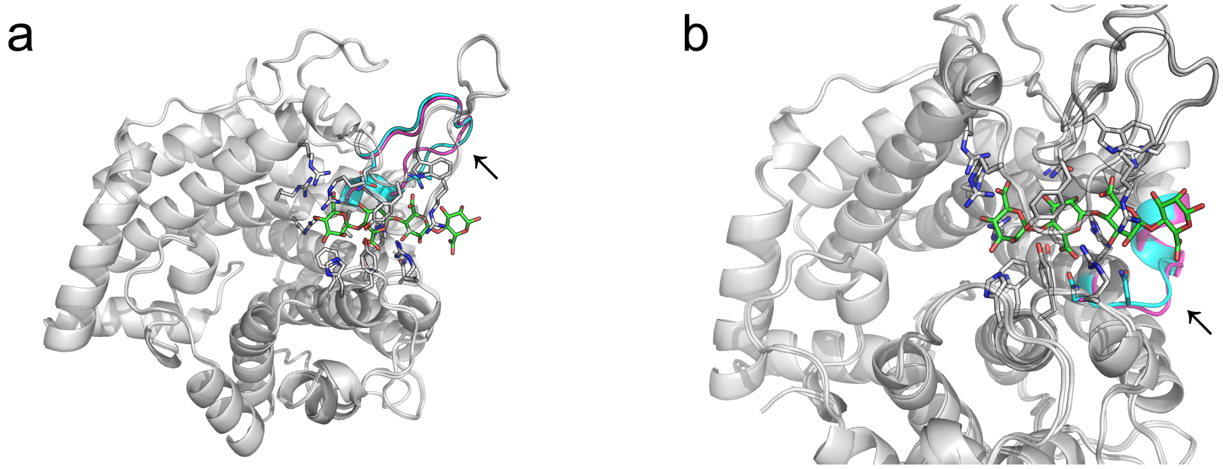

**Supplementary Fig. 12 | Structural disorder induced by mutations.** Loop regions of varants (magenta) change slightly in conformation compared to the same region of WT structure (cyan). **a** Q173N (highlighted region is 173-188. Indicated by arrow), **b** N242A (highlighted region is 235-242, indicated by arrow).

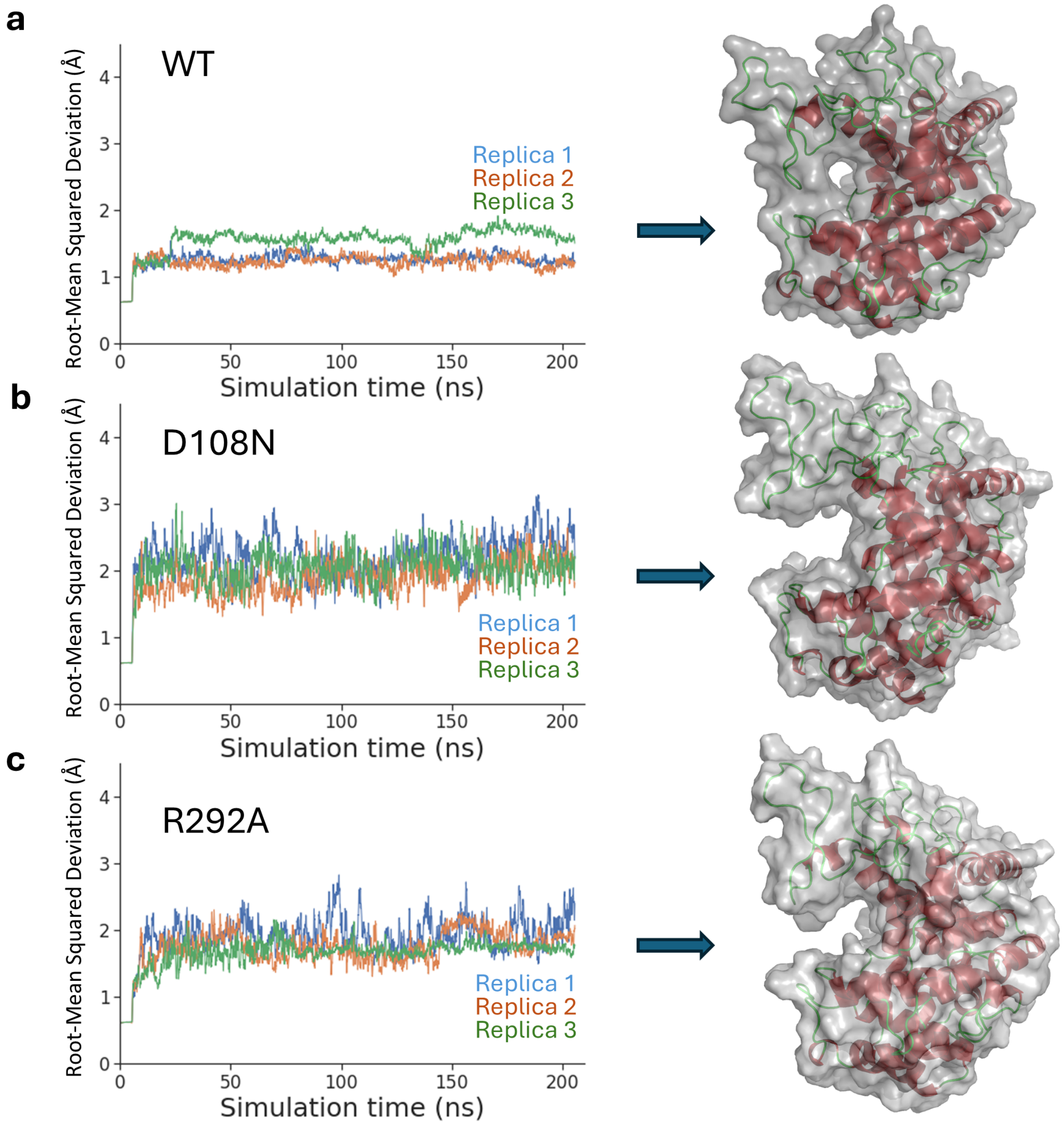

**Supplementary Fig. 13 | Stability analysis of *Bo*PL38 WT (a), D108N (b) and R292A (c) classical MD simulations.** Representative structural conformations after 150 ns is shown on the right of each plot, highlighting the conformational change observed in the active site tunnel.

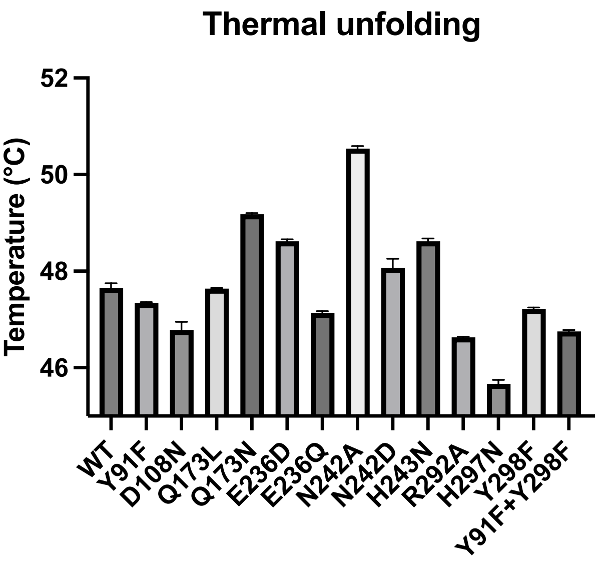

**Supplementary Fig. 14 | Thermal stability analysis of *Bo*PL38 WT and variants.** The thermal stability was assessed using nDSF. This revealed several mutations that seemed to increase the thermal stability. Inspection of the placement of tryptophan residues of the enzyme revealed that the destabilization and loss of tunnel architecture should have no influence on thermal stability readings. Any increase or decrease of thermal stability as monitored using nDSF should be attributed to unfolding processes after the loss of active site integrity.

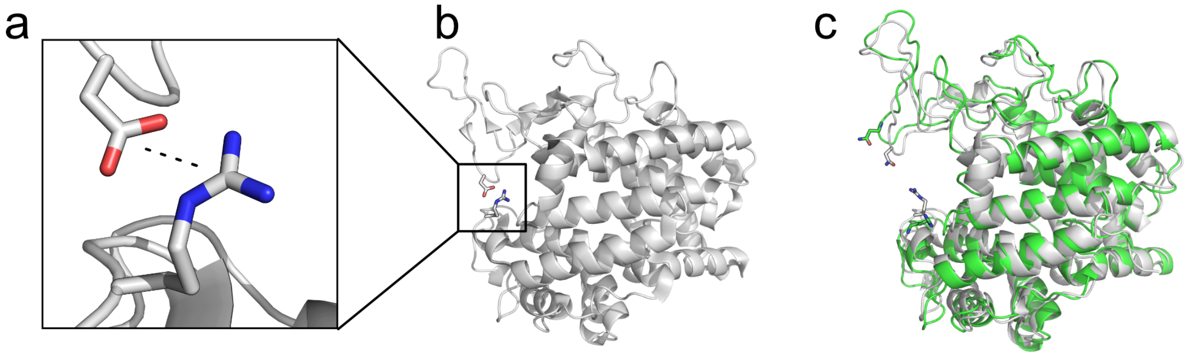

**Supplementary Fig. 15 | Comparing MD simulation with crystal structures. a** A zoom-in on the two residues, D108 and R292, which form a salt bridge in the WT structure of *Bo*PL38 of 3.0 Å. **b** A zoomed-out view of *Bo*PL38, highlighting the salt bridge. **c** Average structural conformations of the MD simulations of D108N and R292A (green) display an opened and unstable active site (see also Supplementary Fig. 13). These conformations are confirmed by the crystal structure of *Bo*PL38-D108N (PDB 9FI0), which adopts the same open conformation (gray) (1.5 Å RMSD to MD simulation snapshot at 150 ns, 2717 atoms) in the absence of the salt bridge.

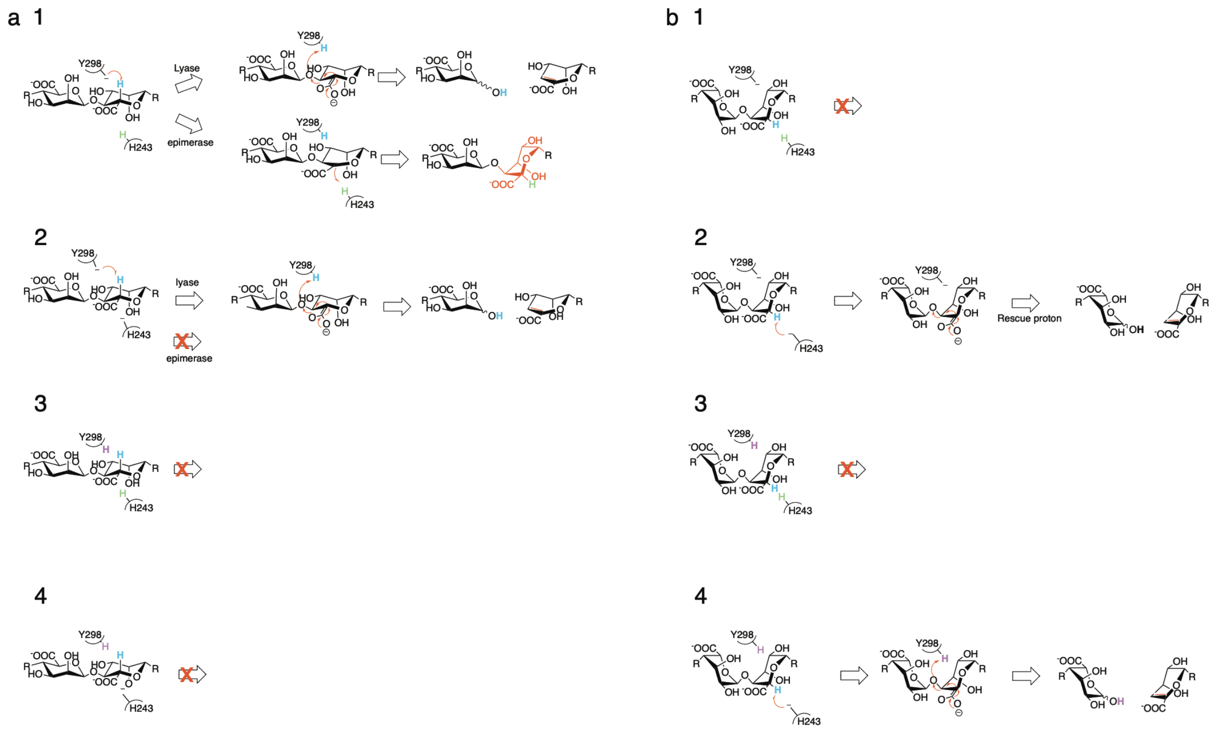

**Supplementary Fig. 16 | Reactions allowed under different protonation conditions of Y298 and H243.** **a** polyM. **b** polyG. **1,** Y298 is deprotonated, H243 is protonated. No reaction will occur on polyG. Lyase or epimerase reactions are possible on polyM. **2**, Both are deprotonated. Both polyG and polyM reactions are possible. No epimerization is possible. **3**, Both are protonated. No reaction. **4**, Y298 is protonated, H243 is deprotonated. This will allow polyG reaction, but not polyM reaction.

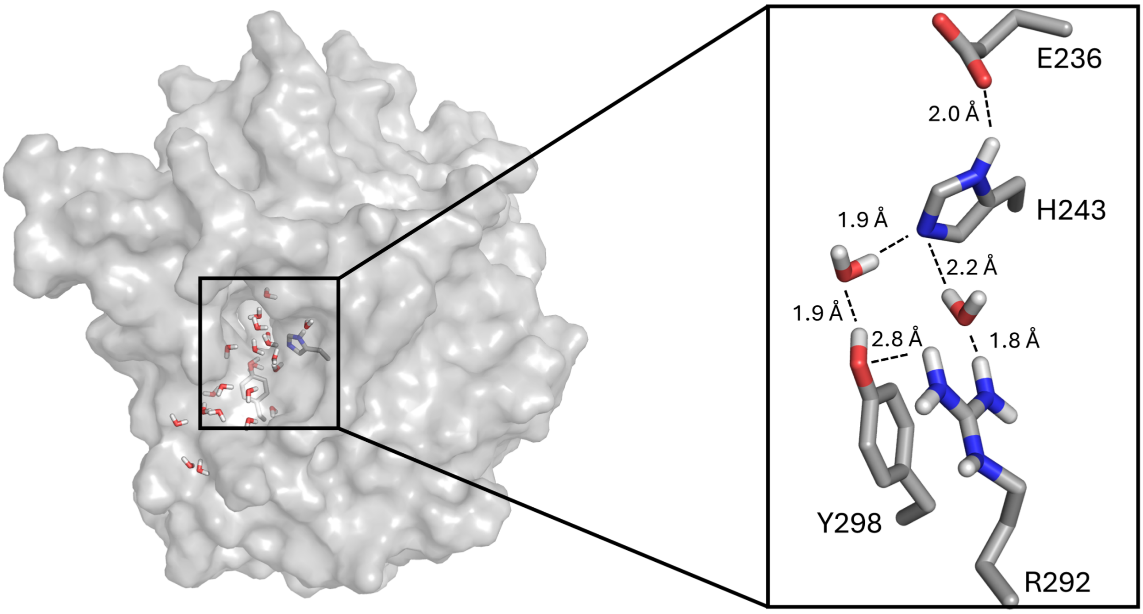

**Supplementary Fig. 17 | Possible water-mediated proton shuttle.** Water molecules observed permeating the tunnel architecture of *Bo*PL38 in MD simulations. The zoom-in illustrates the catalytic site residues, demonstrating their suitability for proton exchange through a water-mediated proton shuttle.

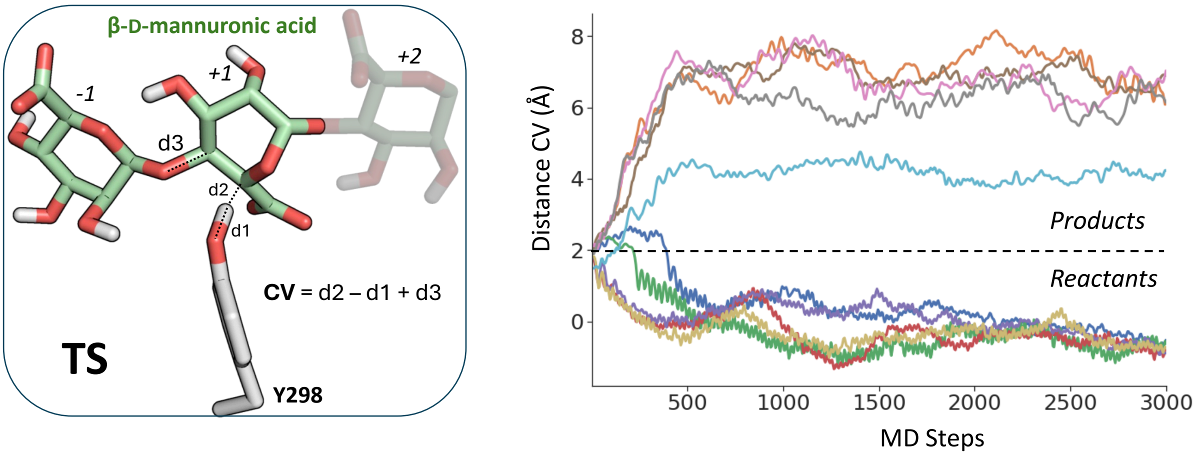

**Supplementary Fig. 18 | Iso-committor analysis of the TS found in the modelled *syn* β-elimination mechanism of *Bo*PL38 on mannuronic acid substrates.** Active site zoom-in of the structure used for the 10 QM/MM runs with different initial velocities is shown on the left. The CV (linear combination of catalytically relevant distances) to follow the evolution is also described. On the right, time evolution of the CV for each of the 10 independent QM/MM MD runs. A dashed line highlights the value of the CV at the initial structure; values larger/smaller than 2 Å indicate simulations evolving towards reactants/products, respectively.

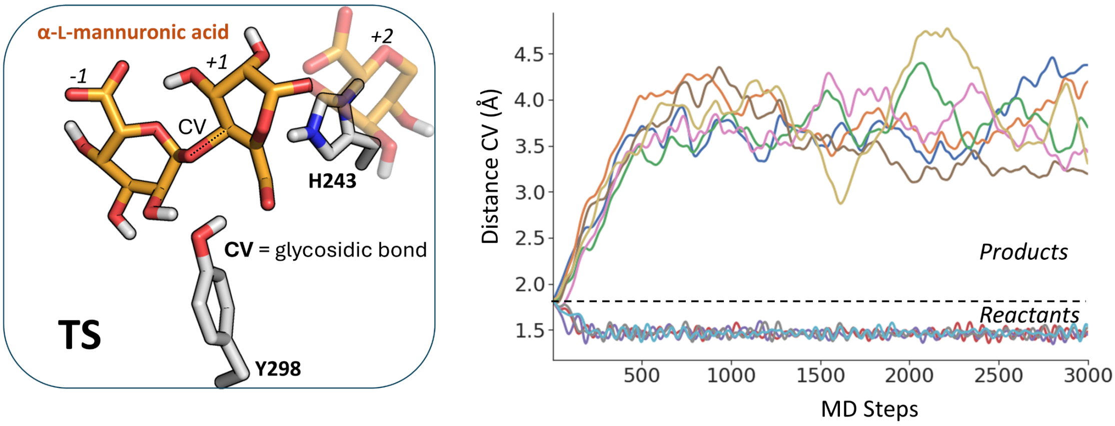

**Supplementary Fig. 19 | Iso-committor analysis of the TS found in the modelled *anti* β-elimination mechanisms of *Bo*PL38 on guluronic acid substrates.** Active site zoom-in of the structure used for the 10 QM/MM runs with different initial velocities are shown on the left. The CV to follow the evolution (glycosidic bond distances) is also described. On the right, time evolution of the CV for each of the 10 independent QM/MM MD runs. A dashed line highlights the value of the CV at the initial structure; values larger/smaller than 1.85 Å indicate simulations evolving towards reactants/products, respectively.

**Supplementary Table 1 | Data collection and refinement statistics for *Bo*PL38-WT and variant structures in complex with alginate oligosaccharides.**

Numbers in parentheses correspond to highest resolution shell.

| *Bo*PL38 | WT-$G_6$ pH 3.5 | WT-$M_6$ pH 3.5 | WT-$(MG)_3$ pH 3.5 | WT-$G_4$ pH 8 | WT-$M_4$ pH 8 | WT-$\Delta M_3$ pH 5 | WT-$\Delta G_3$ pH 5 | D108N-$M_4$ pH 3.5 | H297N-$M_4$ pH 3.5 | N242A-$M_4$ pH 3.5 | Q173N-$G_4$ pH 5 | Q173N-$M_4$ pH 5 | Y298F-$M_4$ pH 3.5 | Y91F-Y298F-$M_4$ pH 3.5 |
|---|---|---|---|---|---|---|---|---|---|---|---|---|---|---|
| **PDB ID** | 9FHT | 9FHU | 9FHV | 9FHW | 9FHX | 9FHY | 9FHZ | 9FI0 | 9FI1 | 9FI2 | 9FI3 | 9FI4 | 9FI5 | 9FI6 |
| **Wavelength** | 0.9999 | 0.9762 | 0.9762 | 0.9762 | 0.8731 | 0.9677 | 0.9677 | 1.0332 | 1.0332 | 1.0332 | 0.9762 | 0.9999 | 0.9724 | 1.0332 |
| **Resolution range** | 42.54 - 2.05<br>(2.07 - 2.05) | 48.93 - 2.09<br>(2.12 - 2.09) | 48.99 - 2.34<br>(2.37 - 2.34) | 62.95 - 2.54<br>(2.57 - 2.54) | 79.08 - 2.11<br>(2.15 - 2.11) | 58.48 - 1.76<br>(1.78 - 1.76) | 52.94 - 1.88<br>(1.9 - 1.88) | 48.49 - 1.86<br>(1.88 - 1.86) | 48.48 - 2.36<br>(2.39 - 2.36) | 49.16 - 2.57<br>(2.6 - 2.57) | 70.34 - 1.99<br>(2.01 - 1.99) | 48.72 - 2.47<br>(2.5 - 2.47) | 48.36 - 2.37<br>(2.4 - 2.37) | 97.05 - 2.18<br>(2.21 - 2.18) |
| **Space group** | *C* 1 2 1 | *C* 1 2 1 | *C* 1 2 1 | *C* 1 2 1 | *C* 1 2 1 | *C* 1 2 1 | *C* 1 2 1 | $P\,2_1\,2_1\,2_1$ | *C* 1 2 1 | *C* 1 2 1 | *C* 1 2 1 | *C* 1 2 1 | *C* 1 2 1 | *C* 1 2 1 |
| **Unit cell** | 197.21<br>88.85<br>146.58<br>90 120.35 90 | 197.84<br>89.01<br>146.82<br>90 120.49 90 | 198.68<br>89.7<br>146.98<br>90 120.56 90 | 196.93<br>89.03<br>145.90<br>90 120.36 90 | 97.33<br>89.36<br>146.51<br>90 120.56 90 | 197.87<br>88.70<br>147.35<br>90 120.39 90 | 198.42<br>88.65<br>147.31<br>90 120.42 90 | 54.74<br>104.49<br>151.49<br>90 90 90 | 197.44<br>89.09<br>146.66<br>90 120.55 90 | 198.57<br>89.15<br>147.48<br>90 120.45 90 | 196.78<br>89.13<br>145.83<br>90 120.38 90 | 197.15<br>88.79<br>146.17<br>90 120.25 90 | 196.68<br>88.87<br>145.09<br>90 120.18 90 | 197.72<br>89.58<br>146.85<br>90 120.39 90 |
| **Total reflections** | 987160<br>(33885) | 255278<br>(6606) | 664656<br>(21574) | 506227<br>(19519) | 832272<br>(41834) | 2258377<br>(46945) | 1236339<br>(24247) | 537456<br>(21719) | 254786<br>(8560) | 333044<br>(11442) | 1043995<br>(34680) | 536013<br>(19559) | 281726<br>(10049) | 803428<br>(27031) |
| **Unique reflections** | 279051<br>(9223) | 250878<br>(6533) | 93431<br>(3000) | 71977<br>(2722) | 124537<br>(6169) | 213920<br>(6126) | 174260<br>(4311) | 73835<br>(2850) | 87904<br>(2917) | 70339<br>(2471) | 148882<br>(4910) | 78192<br>(2755) | 87072<br>(2953) | 115025<br>(3788) |
| **Multiplicity** | 6.9 (7.3) | 6.8 (6.2) | 7.1 (7.2) | 7.0 (7.2) | 6.7 (6.8) | 10.6 (7.7) | 7.1 (5.6) | 7.3 (7.6) | 2.9 (2.9) | 4.7 (4.6) | 7.0 (7.1) | 6.9 (7.1) | 3.2 (3.4) | 7.0 (7.1) |
| **Completeness (%)** | 98.13 (97.25) | 99.37 (92.6) | 99.30 (96.5) | 99.65 (95.92) | 99.90 (100.0) | 98.3 (86.2) | 97.10 (75.9) | 99.92 (100.0) | 97.39<br>(98.48) | 99.12 (90.31) | 96.62 (79.96) | 99.91 (99.93) | 98.94 (99.90) | 100.0 (99.7) |
| **Mean I/sigma(I)** | 11.17 (0.92) | 9.56 (1.32) | 11.8 (0.82) | 6.56 (0.65) | 10.2 (2.1) | 6.6 (0.20) | 6.82 (0.32) | 13.07 (1.66) | 8.05 (1.27) | 7.63 (0.99) | 8.88 (1.14) | 8.73 (1.29) | 7.75 (1.30) | 5.7 (0.40) |
| **Wilson B-factor** | 47.30 | 36.28 | 58.83 | 61.02 | 29.85 | 32.53 | 36.54 | 32.04 | 44.88 | 57.36 | 38.06 | 43.59 | 49.21 | 46.64 |
| **R-merge** | 0.081 (1.891) | 0.094 (0.822) | 0.134 (2.862) | 0.248 (2.997) | 0.137 (0.984) | 0.164 (2.872) | 0.162 (2.649) | 0.084 (1.243) | 0.095(0.823) | 0.125 (1.195) | 0.222 (2.888) | 0.171 (1.396) | 0.100 (0.912) | 0.211 (2.804) |
| **R-meas** | 0.088 (2.036) | 0.102 (0.894) | 0.145 (3.085) | 0.269 (3.233) | 0.148 (1.066) | 0.172 (3.078) | 0.175 (2.909) | 0.091 (1.334) | 0.115(0.995) | 0.141 (1.347) | 0.241 (3.126) | 0.186 (1.506) | 0.120 (1.084) | 0.229 (3.026) |
| **R-pim** | 0.034 (0.751) | 0.039 (0.347) | 0.055 (1.145) | 0.102 (1.204) | 0.057 (0.407) | 0.052 (1.089) | 0.066 (1.178) | 0.033 (0.479) | 0.064(0.553) | 0.064 (0.610) | 0.092 (1.183) | 0.072 (0.562) | 0.066 (0.580) | 0.087 (1.127) |
| **CC1/2** | 0.997 (0.44) | 0.998 (0.720) | 0.995 (0.378) | 0.982 (0.26) | 0.996 (0.677) | 0.996 (0.304) | 0.992 (0.281) | 0.998 (0.729) | 0.995(0.455) | 0.995 (0.367) | 0.988 (0.215) | 0.993 (0.585) | 0.992 (0.494) | 0.972 (0.245) |
| **Reflections used in refinement** | 134497 (4484) | 128726 (3578) | 90774 (2359) | 71746 (2634) | 124507 (4113) | 177043 (97) | 165766 (1387) | 73816 (2849) | 87889(2916) | 70311 (2461) | 143891 (3959) | 78304 (2753) | 87033 (2951) | 109401 (2156) |
| **R-work (%)** | 23.42 | 20.59 | 19.63 | 20.55 | 17.65 | 21.69 | 22.69 | 19.42 | 17.92 | 20.85 | 25 | 20.57 | 17.53 | 22.46 |
| **R-free (%)** | 26.72 | 24.83 | 23.81 | 24.87 | 21.9 | 24.37 | 25.78 | 22.14 | 22.19 | 25.5 | 27.68 | 25 | 21.36 | 26.55 |
| **Number of non-hydrogen atoms** | 13069 | 13615 | 12608 | 12745 | 13367 | 13124 | 12802 | 6677 | 12939 | 12301 | 12549 | 12479 | 12857 | 12884 |
| **macromolecules** | 12208 | 12331 | 12252 | 12202 | 12283 | 12234 | 12215 | 6247 | 12317 | 12243 | 12209 | 12204 | 12223 | 12210 |
| **ligands** | 211 | 231 | 254 | 211 | 196 | 154 | 151 | 0 | 45 | 0 | 10 | 0 | 216 | 226 |
| **solvent** | 650 | 1053 | 102 | 332 | 888 | 736 | 436 | 430 | 577 | 58 | 330 | 275 | 418 | 448 |
| **Protein residues** | 1519 | 1523 | 1520 | 1519 | 1521 | 1520 | 1519 | 762 | 1523 | 1525 | 1520 | 1520 | 1520 | 1522 |
| **RMS(bonds)** | 0.008 | 0.008 | 0.009 | 0.009 | 0.008 | 0.008 | 0.008 | 0.007 | 0.008 | 0.009 | 0.008 | 0.008 | 0.008 | 0.009 |
| **RMS(angles)** | 1.00 | 0.98 | 1.14 | 1.09 | 0.95 | 0.97 | 0.98 | 0.88 | 0.91 | 0.99 | 1.03 | 0.98 | 0.99 | 1.10 |
| **Ramachandran favored (%)** | 97.95 | 97.89 | 97.42 | 97.29 | 97.88 | 98.21 | 98.48 | 98.15 | 97.82 | 96.90 | 97.09 | 96.83 | 97.42 | 97.88 |
| **Ramachandran allowed (%)** | 2.05 | 2.11 | 2.58 | 2.71 | 2.12 | 1.79 | 1.52 | 1.85 | 2.18 | 3.03 | 2.65 | 3.04 | 2.58 | 2.12 |
| **Ramachandran outliers (%)** | 0.00 | 0.00 | 0.00 | 0.00 | 0.00 | 0.00 | 0.00 | 0.00 | 0.00 | 0.07 | 0.26 | 0.13 | 0.00 | 0.00 |
| **Rotamer outliers (%)** | 0.85 | 0.76 | 0.92 | 0.77 | 0.54 | 0.54 | 0.69 | 1.49 | 0.91 | 1.77 | 0.85 | 1.39 | 0.92 | 1.31 |
| **Clashscore** | 5.00 | 3.37 | 5.05 | 6.38 | 2.22 | 3.46 | 3.54 | 5.18 | 3.13 | 5.17 | 5.55 | 4.45 | 3.65 | 5.56 |
| **Average B-factor** | 47.90 | 38.18 | 64.16 | 64.65 | 32.15 | 35.28 | 39.77 | 35.99 | 49.47 | 62.99 | 40.85 | 48.47 | 52.18 | 52.29 |
| **macromolecules** | 47.60 | 37.49 | 63.85 | 64.56 | 31.60 | 35.04 | 39.70 | 35.78 | 49.38 | 63.04 | 40.87 | 48.61 | 51.84 | 51.97 |
| **ligands** | 56.89 | 44.88 | 83.05 | 80.37 | 48.32 | 39.81 | 44.46 | - | 82.09 | - | 96.31 | - | 74.73 | 69.37 |
| **solvent** | 50.49 | 44.83 | 54.81 | 57.97 | 36.19 | 38.30 | 40.23 | 38.95 | 48.75 | 50.99 | 38.36 | 42.24 | 50.30 | 52.31 |

**Supplementary Table 2 | Geometries of bound sugars in *Bo*PL38-$M_4$, -$G_6$ and -$(MG)_2M$ complexes.**

Values separated by slashes are for molecule chains A/B/C/D

| | -2 | -1 | +1 | +2 | +3 | -1 to +3 |
|---|---|---|---|---|---|---|
| RMSD | | | | | | |
| BOUND/RELAXED $M_4$ | | 0.079 Å | 0.18 Å | 0.095 Å | 0.079 Å | 0.53 Å |
| BOUND/RELAXED $G_4$ | | 0.32 Å | 0.29 Å | 0.084 Å | 0.094 Å | 0.88 Å |
| RELAXED $M_4$/$G_4$ | | 0.45 Å | 0.44 Å | 0.46 Å | 0.45 Å | 1.37 Å |
| BOUND $M_4$/$G_4$ | | 0.44 Å | 0.28 Å | 0.50 Å | 0.43 Å | 0.81 Å |
| DISTANCE O1-O4 | | | | | | |
| BOUND $M_4$ | | 5.5 Å | 5.2 Å | 5.4 Å | 5.4 Å | 20.2 Å |
| RELAXED $M_4$ | | 5.7 Å | 5.5 Å | 5.7 Å | 5.5 Å | 20.7 Å |
| BOUND $G_4$ | | 5.2 Å | 5.3 Å | 4.5 Å | 4.7 Å | 18.8 Å |
| RELAXED $G_4$ | | 4.7 Å | 4.7 Å | 4.7 Å | 4.7 Å | 18.5 Å |
| $M_4$ (PRIVATEER) | | | | | | |
| CONFORMATION | | 4C1/4C1/4C1/4C1 | 2H3/4C1/4C1/4C1 | 4C1/4C1/4C1/4C1 | 4C1/4C1/4C1/OE | |
| Q | | 0.6/0.6/0.6/0.6 | 0.4/0.5/0.5/0.5 | 0.6/0.6/0.6/0.6 | 0.5/0.6/0.6/0.5 | |
| PHI | | 358.7/349.7/337.1/284.8 | 161.7/58.6/60.6/99 | 205.4/304.4/348.5/333.5 | 171.4/8.1/60/356.9 | |
| THETA | | 9.5/14.8/18.1/6.7 | 42/6.1/6.3/13.7 | 9.4/4.6/8.3/9.2 | 1.8/12.2/13.2/25.1 | |
| RSCC | | 0.69/0.72/0.66/0.69 | 0.85/0.8/0.81/0.79 | 0.85/0.89/0.75/0.79 | 0.73/0.73/0.58/0.67 | |
| B-FACTOR | | 44.4/42/52.7/53.4 | 34.7/31.9/40.3/48.2 | 32.2/31/41.4/42.6 | 46/44.2/54.6/56.3 | |
| $G_4$ (PRIVATEER) | | | | | | |
| CONFORMATION | | 2SO/2SO/2SO/2SO | 2SO/2SO/2SO/2SO | 1C4/1C4/1C4/1C4 | 1C4/1C4/1C4/1C4 | |
| Q | | 0.7/0.7/0.7/0.7 | 0.6/0.6/0.6/0.7 | 0.6/0.6/0.6/0.6 | 0.5/0.6/0.5/0.6 | |
| PHI | | 152.9/149.8/157.1/154.1 | 143.4/145.6/145.9/148.5 | 296.4/145/143.4/109.1 | 175.8/147/158.5/142.1 | |
| THETA | | 108.2/99.8/105.7/107.3 | 112.3/106.3/110.7/106.1 | 173.6/172.7/173.2/176.7 | 165.9/167.8/164.8/164.5 | |
| RSCC | | 0.7/0.8/0.9/0.7 | 0.8/0.9/0.9/0.8 | 0.8/0.9/0.9/0.9 | 0.8/0.8/0.8/0.8 | |
| B-FACTOR | | 69.8/50.6/51.2/72.1 | 46.9/39/38.9/63.2 | 48.5/39.2/40/59.1 | 71.3/56.5/59.8/76.8 | |
| $(MG)_3$ (PRIVATEER) | | | | | | |
| CONFORMATION | 4C1/4C1/4C1/4C1 | 1C4/1C4/1C4/1C4 | 4C1/4C1/4C1/4C1 | 1C4/1C4/1C4/1C4 | 4C1/4C1/4C1/4C1 | |
| Q | 0.6/0.6/0.6/0.5 | 0.6/0.6/0.6/0.6 | 0.5/0.5/0.5/0.5 | 0.7/0.7/0.6/0.6 | 0.6/0.6/0.6/0.6 | |
| PHI | 291.2/72.7/158.3/193.9 | 279.8/265.4/345.4/300.5 | 80.4/37/108.7/88.5 | 243.9/227.1/258.3/247.6 | 6.2/20.9/318.7/2.2 | |
| THETA | 2.9/2.3/5.6/10.5 | 177.3/179.5/174.4/173.9 | 16.1/12.2/17/14 | 161.9/165/168.6/165.2 | 13.3/22.4/3.9/8.5 | |
| RSCC | 0.7/0.6/0.7/0.7 | 0.9/0.8/0.9/0.9 | 0.9/0.8/0.9/0.9 | 0.8/0.8/0.8/0.8 | 0.6/0.7/0.4/0.5 | |
| B-FACTOR | 94.4/104.9/98.2/108.3 | 65.5/70.9/69.8/71 | 56.7/61.6/66.1/60.8 | 76.8/74.7/84.2/79.6 | 112.6/115.6/117.5/118.7 | |

**Supplementary Table 3 |** Assignment of $^1$H and $^{13}$C chemical shifts of Δ-GG trimer. The roman numbers (I-IV) indicate the residue position from the reducing end. The numbers (1-6) indicate the ring carbon numbers.

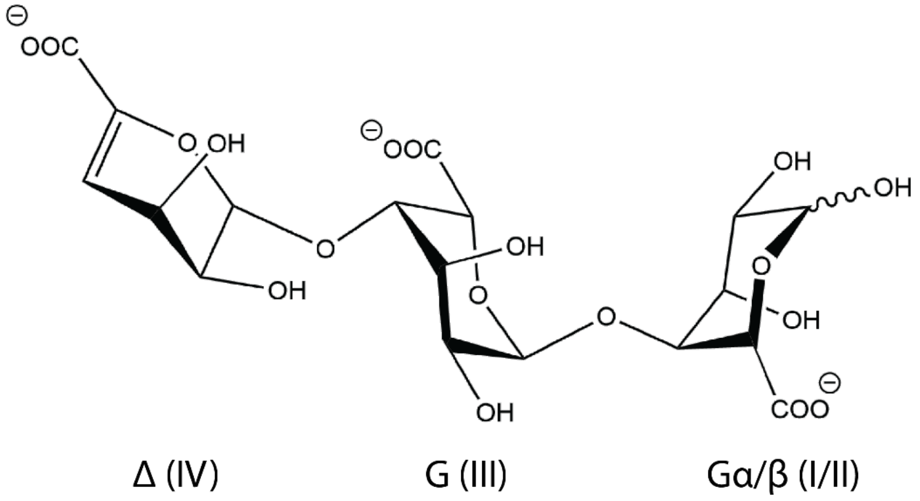

| | H-1; C-1 | H-2; C-2 | H-3; C-3 | H-4; C-4 | H-5; C-5 | C-6 |
|---|---|---|---|---|---|---|
| **Gα I** | 5.19; 95.8 | 3.90; 67.1 | 4.06; 72.7 | 4.12; 83.0 | 4.06; 72.7 | 178.0 |
| **Gβ II** | 4.85; 96.0 | 3.59; 71.8 | 4.09; 72.8 | 4.01; 83.0 | 4.38; 76.1 | 178.0 |
| **G III** | 4.99; 103.4 | 3.84; 67.7 | 4.07; 71.7 | 4.22; 82.5 | 4.45; 69.8 | 178.2 |
| **Δ IV** | 5.16; 103.2 | 3.89; 69.6 | 4.32; 65.4 | 5.82; 110.5 | -; 147.5 | 171.8 |

**Supplementary Table 4 |** Assignment of $^{1}H$ and $^{13}C$ chemical shifts of Δ-MM trimer. The roman numbers (I-V) indicate the residue position from the reducing end. The numbers (1-6) indicate the ring carbon numbers.

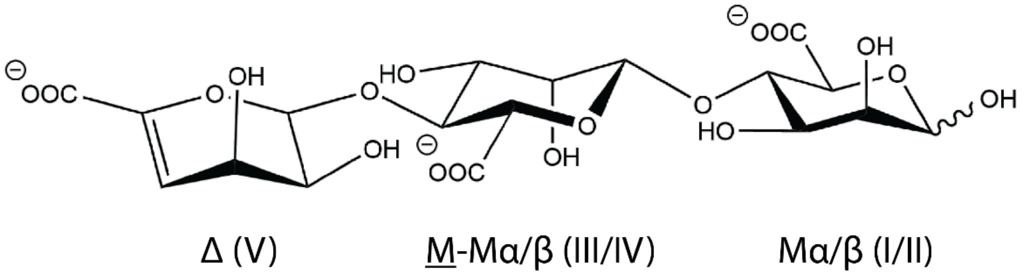

| | H-1; C-1 | H-2; C-2 | H-3; C-3 | H-4; C-4 | H-5; C-5 | C-6 |
|---|---|---|---|---|---|---|
| **Mα I** | 5.19; 96.2 | 3.91; 72.7 | 3.95; 71.7 | 3.99; 81.1 | 4.12; 75.4 | 178.8 |
| **Mβ II** | 4.89; 96.4 | 3.94; 73.3 | 3.79; 74.2 | 3.87; 80.9 | 3.71; 78.9 | 178.8 |
| **M-Mα III** | 4.66; 102.5 | 4.01; 73.1 | 3.74; 74.1 | 3.93; 80.8 | 3.75; 78.5 | 178.1 |
| **M-Mβ IV** | 4.62; 102.7 | 4.01; 73.1 | 3.74; 74.1 | 3.93; 80.8 | 3.75; 78.5 | 178.1 |
| **Δ V** | 5.09; 102.8 | 3.94; 69.4 | 4.42; 66.3 | 5.72; 110.2 | -; 147.9 | 172.0 |

**Supplementary Table 5 |** Assignment of $^{1}H$ and $^{13}C$ chemical shifts of Δ-$G_{\beta}$ dimer. The roman numbers (I-II) indicate the residue position from the reducing end. The numbers (1-6) indicate the ring carbon numbers.

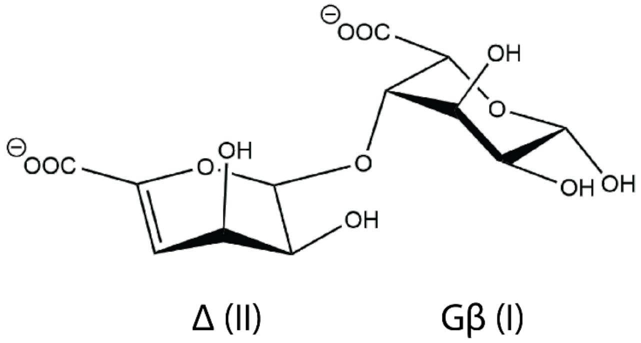

| | H-1; C-1 | H-2; C-2 | H-3; C-3 | H-4; C-4 | H-5; C-5 | C-6 |
|---|---|---|---|---|---|---|
| **Gβ I** | 4.86; 96.0 | 3.55; 71.5 | 4.41; 76.0 | 4.15; 82.4 | 4.18; 72.6 | 178.0 |
| **Δ II** | 5.21; 103.0 | 3.93; 69.6 | 4.31; 65.0 | 5.92; 110.5 | -; 147.2 | 172.0 |

**Supplementary Table 6 |** Assignment of $^{1}H$ and $^{13}C$ chemical shifts of Δ-$M_{\alpha}$ dimer. The roman numbers (I-II) indicate the residue position from the reducing end. The numbers (1-6) indicate the ring carbon numbers.

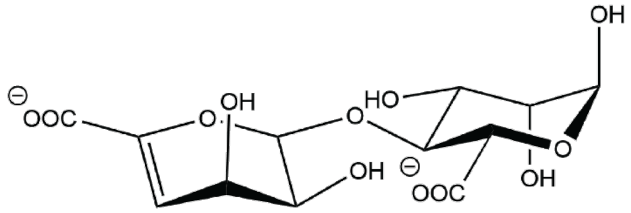

Δ (II) Mα (I)

| | H-1; C-1 | H-2; C-2 | H-3; C-3 | H-4; C-4 | H-5; C-5 | C-6 |
|---|---|---|---|---|---|---|
| **Mα I** | 5.24; 96.0 | 4.10; 71.8 | 4.20; 75.8 | 4.10; 81.3 | N.A.; N.A. | 178.7 |
| **Δ II** | 5.17; 102.5 | 3.97; 69.6 | 4.43; 66.2 | 5.80; 110.4 | -; 147.6 | 172.1 |

**Supplementary Table 7 |** Assignment of $^1$H and $^{13}$C chemical shifts of internal residues: M residue in polyMG (-<u>M</u>G-), G residue in polyMG (-<u>G</u>M-), G residue in polyG (-<u>G</u>G-), and M residue in polyM (-<u>M</u>M-). The numbers (1-6) indicate the ring carbon numbers.

| | **H-1; C-1** | **H-2; C-2** | **H-3; C-3** | **H-4; C-4** | **H-5; C-5** | **C-6** |
|---|---|---|---|---|---|---|
| **-<u>M</u>G-** | 4.70;104.3 | 3.91;73.7 | 3.74;74.3 | 3.77;80.1 | 3.82;78.5 | UA |
| **-<u>G</u>M-** | 5.02;102.4 | 3.97;67.6 | 4.18;72.2 | 4.21;82.9 | 4.80;70.3 | UA |
| **-<u>G</u>G-** | 5.05;103.6 | 3.91;67.9 | 4.01;71.9 | 4.12;82.9 | 4.47;70.0 | UA |
| **-<u>M</u>M-** | 4.65;102.8 | 4.04;72.8 | 3.76;74.0 | 3.89;80.7 | 3.75;78.7 | UA |

**Supplementary Table 8 |** Overview over all time-resolved NMR reactions run with *Bo*PL38 WT and the different substrates showing their reaction and acquisition parameters.

| Substrate | *Bo*PL38 (nM) | Substrate (mg ml$^{-1}$) | Time between scans (min) | Total exp. duration |
|---|---|---|---|---|
| **PolyM** | 412 | 9.30 | 5 | 13 h 20 min |
| **OligoG** | 845 | 7.71 | 5 | 13 h 20 min |
| **PolyMG** | 932 | 8.57 | 5 | 13 h 21 min |
| **$^{13}$C-1 PolyM** | 558 | 8.50 | 5 | 15 h 1 min |

**Supplementary Table 9 |** Overview over all time-resolved NMR reactions run of *Bo*PL38 variants and their reaction and acquisition parameters. Substrate for all reactions is $^{13}$C-1 labelled polyM DP 70–100.

| Variant | Enzyme (μM) | Substrate (mg ml$^{-1}$) | Time between scans (min) | Total exp. duration |
|---|---|---|---|---|
| **Y91F** | 6.05 | 8.35 | 10 | 10 h 40 min |
| **Y91F+Y298F** | 5.64 | 8.98 | 5 | 10 h 1 min |
| **D108N** | 5.49 | 7.57 | 5 | 10 h 1 min |
| **Q173L** | 6.04 | 8.63 | 10 | 10 h 40 min |
| **Q173N** | 6.95 | 8.63 | 10 | 10 h 40 min |
| **E236D** | 5.51 | 7.19 | 5 | 15 h 0 min |
| **E235Q** | 5.49 | 7.75 | 5 | 15 h 0 min |
| **N242A** | 5.60 | 7.90 | 5 | 6 h 30 min |
| **N242D** | 5.59 | 8.00 | 5 | 15 h 0 min |
| **H243N** | 3.73 | 8.33 | 10 | 10 h 40 min |
| **R292A** | 5.49 | 8.53 | 10 | 10 h 40 min |
| **H297N** | 3.65 | 8.33 | 10 | 10 h 40 min |
| **Y298F** | 3.98 | 8.80 | 5 | 12 h 30 min |
